# Defensive fungal symbiosis on insect hindlegs

**DOI:** 10.1101/2024.03.25.586038

**Authors:** Takanori Nishino, Hiromi Mukai, Minoru Moriyama, Takahiro Hosokawa, Masahiko Tanahashi, Shuji Tachikawa, Naruo Nikoh, Ryuichi Koga, Takema Fukatsu

## Abstract

Tympanal organs as “insect ears” have evolved repeatedly. Dinidorid stinkbugs were reported to possess a conspicuous tympanal organ on female’s hindlegs. Here we report an unexpected discovery that the stinkbug’s “tympanal organ” is actually a novel symbiotic organ. The stinkbug’s “tympanum” is not membranous but a porous cuticle, where each pore connects to glandular secretory cells. In reproductive females, the hindleg organ is covered with fungal hyphae growing out of the pores. Upon oviposition, the females skillfully transfer the fungi from the organ to the eggs. The eggs are quickly covered with hyphae and physically protected against wasp parasitism. The fungi are mostly benign Cordycipitaceae entomopathogens and show considerable diversity among insect individuals and populations, indicating environmental acquisition of specific fungal associates. These results uncover a novel external fungal symbiosis in which host’s elaborate morphological, physiological and behavioral specializations underpin the selective recruitment of benign entomopathogens for a defensive purpose.

## Main text

Tympanal organs are the external auditory structures that have repeatedly evolved in diverse insects. Crickets, moths and cicadas have tympanal ears on their forelegs, thorax and abdomen, respectively, by which they perceive courtship songs and recognize approaching enemies (1,2). Not only musical cicadas but also other hemipteran insects, including diverse stinkbugs, utilize either air-borne sounds or substrate-borne vibrations in their mating behaviors (3,4). The Dinidoridae is a small stinkbug family embracing some 100 species representing 16 genera (5–7). Morphological studies reported that dinidorid stinkbugs generally develop a conspicuous tympanal organ specifically on female’s hind tibiae, which was speculated to be used for courtship communications (8). Thus far, however, no detailed studies have been conducted on this minor group of stinkbugs. How do dinidorid females listen to male’s song or dance using their hindlegs? To address this question, we investigated the Japanese dinidorid stinkbug *Megymenum gracilicorne*, and discovered that, unexpectedly, the stinkbug’s “tympanal organ” is not an auditory organ but a previously unknown type of symbiotic organ.

## Results and Discussion

### Peculiar traits of fungus-growing “tympanal organ” on hindlegs

*M. gracilicorne* lives on wild cucurbitaceous plants and sometimes infests cultivated cucumbers and pumpkins (9) (Fig. 1a-c). As previously reported for over 30 dinidorid species (8), *M. gracilicorne* exhibits a conspicuous sexual dimorphism in the hindlegs. While the hind tibiae of adult males are normal in shape (Fig. 1d-f), the hind tibiae of adult females are widened and concave in the middle, where a flat oval area of “tympanum” develops at the center (Fig. 1g, h). However, close morphological inspection uncovered that structural features of the female-specific hindleg organ are atypical of insect’s tympanal ears in that the “tympanum” was not membranous but a rigid porous cuticle (Fig. 1h, i). We found that, in reproductively mature females, the “tympanum” was covered with wooly white materials (Fig. 1b, c, j), which looked like fungal hyphae massively growing out of the pores (Fig. 1k, l). Cultivation and microscopic observation of the white materials confirmed that they are certainly filamentous fungi, stretching hyphae and forming conidia (Fig. S1).

**Fig. 1.**
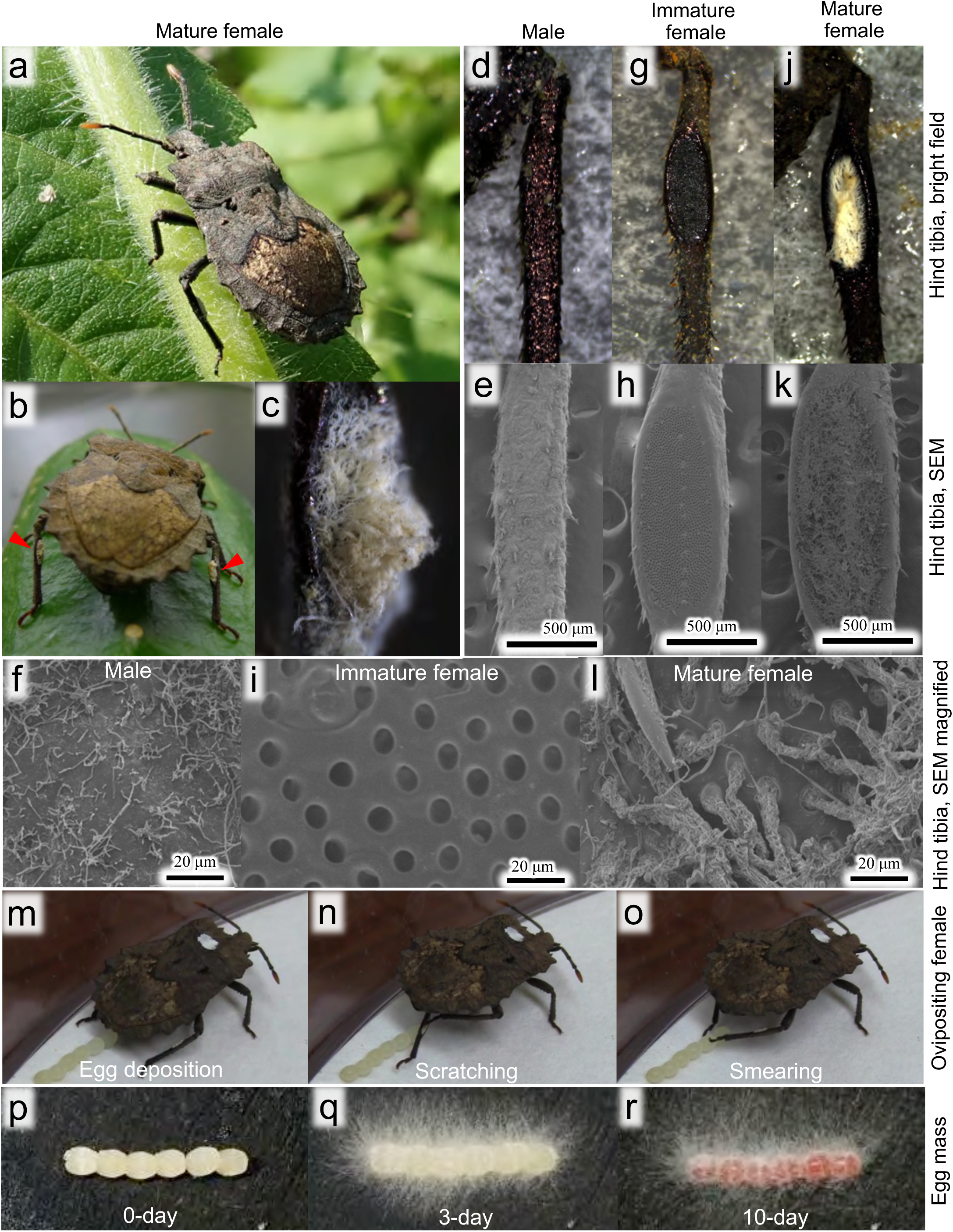
Morphology and behavior of *M. gracilicorne*. (**a**) An adult female in the field on *Sicyos angulatus*. (**b**) Back view of an adult female reared on cucumber, whose hindleg organs are covered with fungal hyphae (arrowheads). (**c**) Magnified image of the hindleg organ. (**d-l**) Morphology of hind tibia of adult male (**d-f**), immature adult female (**g-i**) and mature adult female (**j-l**). (**d, g, i**) Bright field images. (**e, h, k**) SEM images. (**f, i, l**) Magnified SEM images. (**m-o**) Peculiar behavior of egg-laying adult female. After laying each egg (**m**), the female rhythmically scratches the hyphae-covered hindleg organ with tarsal claws of the opposite hindleg (**n**) and then rubs the egg surface with the claws in a skillful manner (**o**). Also see Movie S1. (**p-r**) Egg mass covered with fungal hyphae, 0-day (**p**), 3-day (**q**) and 10-day (**r**) after oviposition. Though densely covered with fungal hyphae, the eggs normally exhibit high hatch rates, typically over 80%.

### Female’s behavior for fungus transfer from hindleg organ to eggs

Strikingly, we observed a peculiar behavior of the reproductive adult females during oviposition. The gravid females laid eggs in a row, and when each egg was deposited, the females rhythmically scratched the hyphae-covered hindleg organ with tarsal claws of the opposite hindleg and then rubbed the egg surface with the claws in a skillful manner, thereby smearing the fungi onto the eggs (Fig. 1m-o; Movie S1). Within a few days, fungal hyphae quickly grew and covered the entire egg mass (Fig. 1p-r). Upon hatching, the hyphae attached to the body surface of newborn nymphs (Fig. S2; Movie S2), but the fungi were subsequently lost as the nymphs molted and grew, being not maintained on the insects.

### Diversity and specificity of symbiotic fungi

In order to identify the fungi growing on the hindleg organs and the eggs, we collected reproductive adult females from different localities, each female was reared in isolation and allowed to lay two egg masses consecutively, and the hind tibiae, the first egg mass and the second egg mass were sampled and subjected to fungal cultivation (Table S1; Fig. S3). In total, over 600 fungal isolates were obtained and subjected to sequencing of ribosomal internal transcribed spacer (ITS) region (Table S1). We also performed amplicon sequencing analysis of the fungal ITS region using the DNA samples extracted from the hind tibiae and the eggs (Table S2). Molecular phylogenetic analyses based on these sequence data consistently revealed that (i) the majority of the fungi were placed within the class Sordariomycetes of the Ascomycota, (ii) the majority were classified to the family Cordycipitaceae mainly consisting of entomopathogenic fungi such as *Cordyceps*, *Beauveria* and others (10,11), and (iii) the majority represented several specific fungal lineages such as *Lecanicillium*, *Simplicillium* and *Akanthomyces* (12–14) (Figs. S4 and S5; Note S1).

### Fungal diversity and specificity in natural host populations

Figure 2 shows the fungal compositions on the hindlegs, the first eggs and the second eggs of field-collected reproductive females of *M. gracilicorne* originated from different localities and years, based on the ITS amplicon sequencing data. The following patterns were observed: (i) each female generally hosted multiple fungal species, reflecting the external open configuration of the symbiotic system vulnerable to microbial contamination; (ii) the fungal compositions on the maternal hindlegs were generally similar to the fungal compositions on the eggs, confirming vertical transmission of the fungal associates by the maternal smearing behavior; (iii) within the same collection locality, the fungal compositions were often markedly different between the insects, highlighting intra-populational diversity of the fungal associates; (iv) the predominant fungal associates may be different among the collection localities, illustrating inter-populational diversity of the fungal associates; and (v) within the same collection locality, the predominant fungal associates may be different across different years. These patterns looked quite strange – despite the fidelity of mother-offspring vertical transmission, the fungal associates exhibit remarkable diversity and variability within and between the populations of *M. gracilicorne*.

**Fig. 2.**
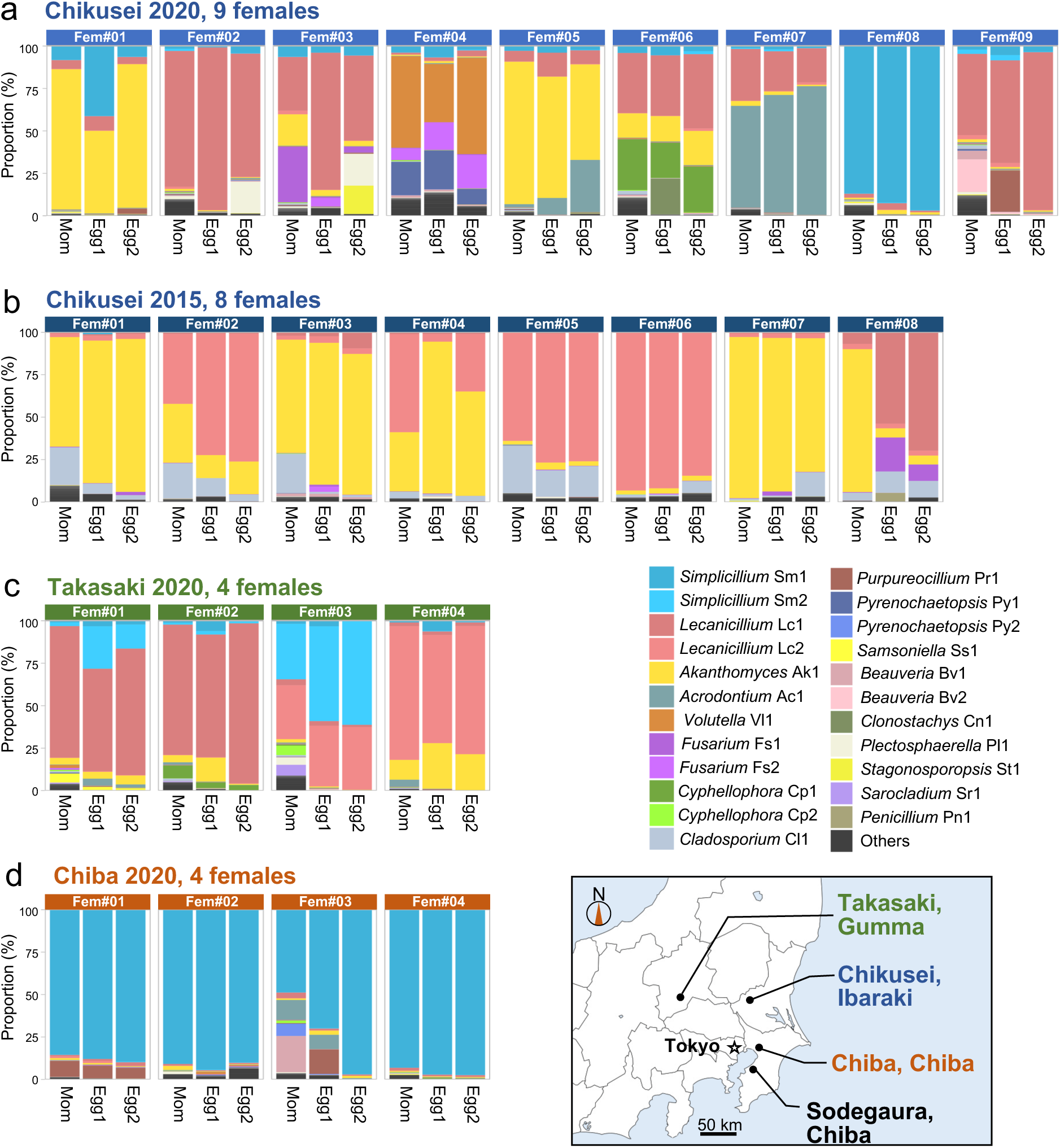
Fungal microbiota on hindleg organs and eggs of field-collected reproductive adult females of *M. gracilicorne*. (**a**) Nine females collected at Chikusei in 2020. (**b**) Eight females collected at Chikusei in 2015. (**c**) Four females collected at Takasaki in 2020. (**d**) Four females collected at Chiba in 2020. Fungal ITS amplicon-seq data from the mother’s hindlegs (Mom), the first egg mass (Egg1) and the second egg mass (Egg2) are shown by colored bar graphs. For detail, also see Fig. S3.

### Vertical transmission, loss, and environmental re-acquisition of symbiotic fungi

These results indicate that *M. gracilicorne* transmits the hindleg-associated fungi to eggs vertically, but the fungi are lost during the nymphal development, and the fungi are newly acquired by adult females from the environment every generation. Such a symbiont transmission trajectory is, to our knowledge, exceptional among insect-microbe and other symbiotic systems (15–17). Why the symbiotic fungi passed to offspring by mother’s specialized morphology and skillful behavior have to be lost and regained is an enigma.

### Structure, cytology, and gene expression of hindleg organ

Another enigma is how the specific fungal strains, either *Lecanicillium*, *Simplicillium* or *Akanthomyces*, can be selectively picked up by the female’s hindleg organ from a myriad of microbes in the environment. To address this question, we investigated the female’s hindleg organ in detail. Histologically, the “tympanum” was a thick and sclerotized porous cuticle, and the levels of thickness and sclerotization were more conspicuous in mature females than in immature females (Fig. 3a-d). Beneath the “tympanum”, no neuronal elements like chordotonal organs, the stretch receptors typically associated with auditory and vibratory sense organs of insects (3,4), were seen, but the thick cuticle was lined with a well-developed epidermal cell layer (Fig. 3a-d). Notably, the epidermal cells intruded into the cuticular layer and connected to the bottom of the cuticular pores, constituting secretory epidermal columns (Fig. 3b, d). In mature females, the epidermal columns and the cuticular pores exhibited dense signals of polysaccharide staining, indicating high secretion activities of the epidermal cells and massive fungal proliferation in the cuticular pores (Fig. 3c, d), which were also verified by TEM observations (Fig. 3e-h). Comparative transcriptomic analyses of midlegs and hindlegs of immature and mature adult insects of *M. gracilicorne*, from which fungus-derived and other presumably contaminating genes were removed based on sequence similarity (Tables S3 and S4), revealed that the hindlegs of mature adult females exhibit a distinct gene expression pattern (Fig. S6). Differential gene expression analyses identified 550 genes highly and preferentially expressed in the hindlegs of mature adult females (Tables S5 and S6). Functional term enrichment analysis revealed that genes with transporter activities were augmented in the hindleg organ, which was consistent with its secretory role. The top highly expressed gene encoded a Takeout family protein, which is a small secretory protein likely functioning as a carrier of hydrophobic substances (18). The other highly expressed genes included UDP-glucuronosyltransferases and glucuronate pathway genes that presumably mobilize lipophilic materials, a secretory cuticular glycoprotein possibly involved in the formation of the cuticular structure, some immune-related regulators in the Toll signaling pathway, and others (Table S6; Note S2).

**Fig. 3.**
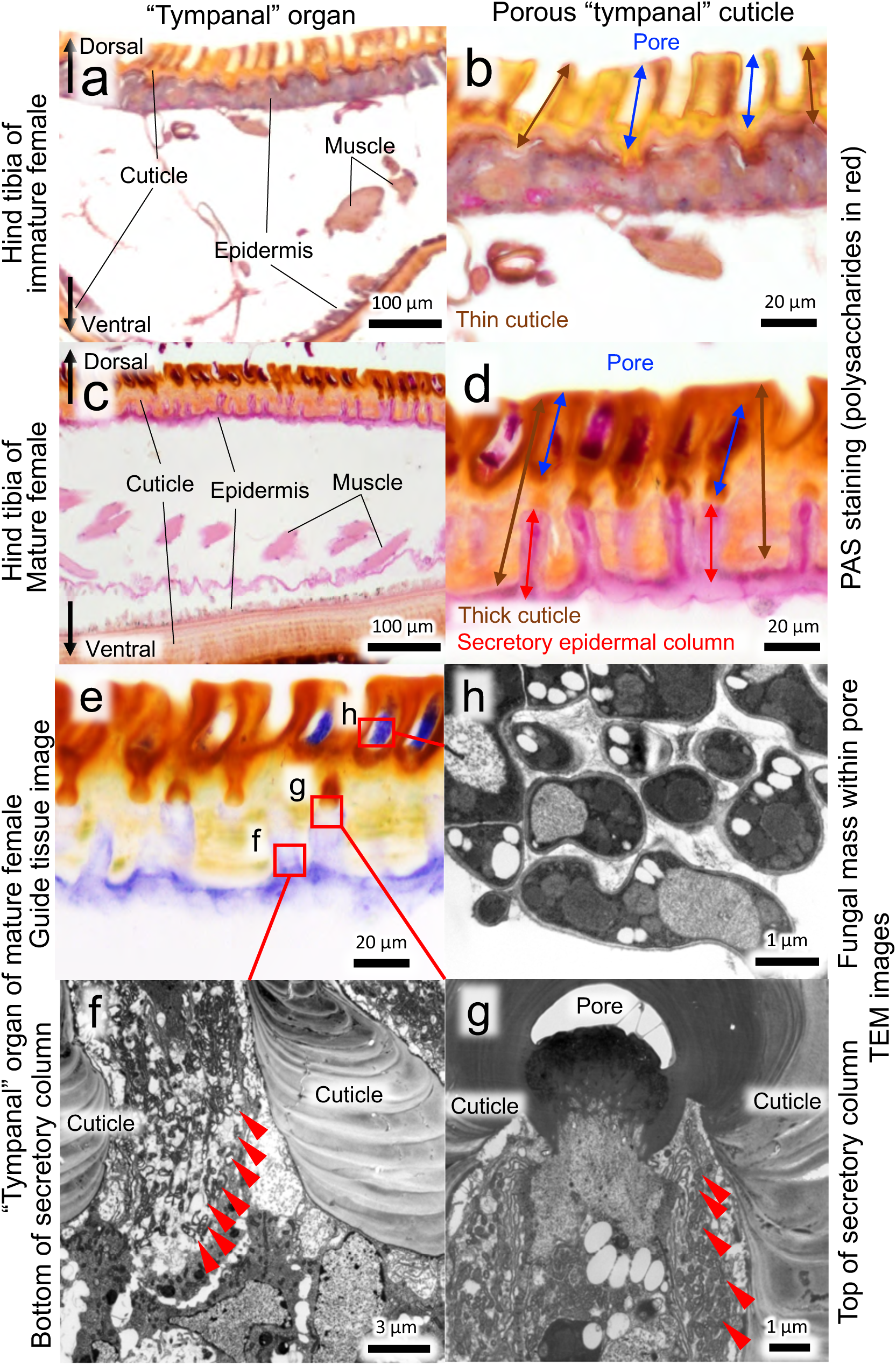
Histology and fine structure of female-specific hindleg organ of *M. gracilicorne*. (**a-d**) Tissue section images of the hindleg organs subjected to PAS staining, by which sugar-rich elements such as polysaccharides, glycoproteins, mucous secretion and fungal cell wall are visualized in red. (**a, b**) In immature female, the “tympanal” cuticle is thinner, the epidermal cells are stained in pale pink, and the cuticular pores are empty. (**c, d**) In mature female, the “tympanal” cuticle is thicker and sclerotized, the epidermal cells are deeply stained in red, and the cuticular pores are full of fungal hyphae. Note that no neuronal elements like chordotonal organs are found beneath the “tympanal” cuticle. Also note that the epidermal cells form secretory columns leading to the bottom of the cuticular pores. (**e**) A tissue section image of the “tympanal” cuticle of mature female for guiding the histological locations corresponding to TEM images (**f-h**) shown by red rectangles. (**f**) TEM image of the bottom of a secretory column. The cytoplasm of secretory epidermal cells full of granules, vacuoles and membranous elements is seen between thick cuticles. (**g**) TEM image of the top of a secretory column. The cytoplasm full of secretory granules and membranous elements ends up with the narrow cuticular canals at the bottom of the cuticular pore. Arrowheads highlight secretory granules. (**h**) TEM image of the content of a cuticular pore. Densely packed fungal hyphae are seen.

### Hindleg organ as external symbiotic organ for hosting specific fungi

These results unequivocally rejected the conventional interpretation that the stinkbug’s female-specific hindleg organs are tympanal ears, which was based merely on superficial morphological resemblance (8). The “tympanum” is neither a vibratory membrane nor associated with sensory neurons, but is a sclerotized cuticle with numerous pores whose bottom is connected to secretory cells. The morphological and cytological features of the pores are reminiscent of the glandular mycangia, the exoskeletal cavities lined with glandular cells for fostering specific symbiotic fungi as known from ambrosia beetles (19,20). Similar exoskeletal cavities with glandular elements for hosting symbiotic actinobacteria are reported from leaf-cutter ants and digger wasps (21,22). The peculiarity in *M. gracilicorne* is the large number and high density of the mycangial units concentrated in the specialized region on the hindlegs. While ambrosia beetles generally have only a few mycangia (23), each hindleg organ of *M. gracilicorne* bears about 2,000 fungus-growing pores (mean ± SD = 2,118 ± 100; n = 11), totaling over 4,000 glandular mycangia per adult female. We conclude that the hindleg organ is a previously unknown type of external symbiotic organ consisting of highly concentrated glandular mycangia for hosting specific fungal associates.

### Egg surface structure, fungal growth and proteinaceous secretion

Following the maternal fungus-smearing behavior upon oviposition (Fig. 1m-o; Movie S1), the egg surface is soon covered with fungal hyphae (Fig. 1p-q). What support the massive and quick fungal growth on the eggs? SEM and sectioning observations revealed that the egg surface is covered with a polysaccharide-rich layer and the fungal hyphae grow presumably by consuming the secretion layer (Fig. S7a-f). The secretion layer was already observed on the surface of mature eggs in the female’s ovary (Fig. S8a). The secretion layer was solubilized when disulfide bond was reduced in the presence of detergent (Fig. S8b). From newly laid eggs, the secretion layer was extracted (Fig. S7g-k) and subjected to proteomic analysis (Table S7). An outstandingly abundant secretion protein with an odorant-binding protein (OBP)-like motif was identified. The other detected proteins were β-glucuronidase, probable antibacterial peptide, soma ferritin, and others (Table S7; Note S2). A recent work reported that, for vertical transmission upon oviposition, stinkbugs of the family Plataspidae produce symbiont-containing “capsules”, in which a predominant secretion protein with an OBP domain, called PMDP, embeds and protects the fragile symbiont cells outside the host (24). However, the abundant OBP-like motif protein of *M. gracilicorne* was phylogenetically not related to PMDP. On account of the quantitative predominance, it is conceivable, although speculative, that the protein might contribute to the fungal proliferation on the egg surface at least to some extent.

### Low to moderate pathogenicity of hindleg-associated fungi

The Cordycipitaceae is famous as a fungal group containing many potent entomopathogens such as *Cordyceps* and *Beauveria* (10,11). Keeping such entomopathogens on the hindlegs and eggs seems dangerous for the host insect. However, we found that the major hindleg-associated fungal strains may be not so dangerous for *M. gracilicorne*. Topical application of conidia, by dipping the adult insects into conidia suspension of *Simplicillium* Sm1, *Lecanicillium* Lc2 or *Akanthomyces* Ak1, did not kill the insects (Fig. S9a). When fungal conidia were injected into hemocoel, a representative pathogen *Beauveria* promptly killed all the injected insects. By contrast, the hindleg-derived *Simplicillium* Sm1 and *Lecanicillium* Lc2 showed very low mortality of the injected insects whereas *Akanthomyces* Ak1 exhibited some pathogenicity though lower than *Beauveria* (Fig. S9b). From these results, we thought that *M. gracilicorne* might have developed resistance to these entomopathogens during the symbiotic association. However, this idea turned out to be inappropriate when the same experiments were conducted using the pentatomid stinkbug *Plautia stali* with no relationship to these fungi. Similarly, conidia injection experiments revealed that *Beauveria* killed all the injected insects, *Simplicillium* Sm1 and *Lecanicillium* Lc2 killed few insects, and *Akanthomyces* Ak1 killed some insects, whereas topical application of conidia did not kill the insects (Fig. S9c, d). These results suggest that adult females of *M. gracilicorne* preferentially pick up relatively benign entomopathogens, *Simplicillium* or *Lecanicillium* (12–14,25,26), from environmental sources and selectively maintain them on the hindleg organ (Note S3).

### Defensive symbiosis against microbial pathogens?

In such insects as leaf cutter ants (27,28), digger wasps (29,30), bark beetles (31), darkling beetles (32,33), leaf rolling weevils (34), leaf beetles (35) and others, external symbiotic actinobacteria, proteobacteria or ascomycetous fungi were reported to produce potent antimicrobial substances, thereby chemically protecting their host insects against microbial parasites and pathogens (36–38). Hence, we first suspected that the symbiotic fungi of *M. gracilicorne* might also play similar biological roles. Since there was no available information about microbial parasites and pathogens of *M. gracilicorne*, we attempted co-culture of the hindleg-derived *Simplicillium*, *Lecanicillium* and *Akanthomyces* with several bacteria (*Escherichia coli*, *Pantoea dispersa*, *Bacillus subtilis*, etc.) on the same agar plates, but conspicuous suppressive effects were not observed. The RNA sequencing data of the hindlegs of mature adult females contained fungal gene expression data, but no conspicuous expression of antibiotic-related genes like polyketide synthases was detected (Table S8; Note S4). Of course, the possibility that the hindleg-associated fungi are effective against hitherto unidentified microbial enemies of *M. gracilicorne* cannot be excluded. On the other hand, we discovered the relevance of the symbiotic fungi to a non-microbial natural enemy of *M. gracilicorne*.

### Defense against wasp parasitism by egg-covering fungal hyphae

By inspecting field-collected eggs of *M. gracilicorne*, we noticed that the natural eggs frequently suffer wasp parasitism. While newly laid eggs were white in color (Fig. 4a), developing fertilized eggs became reddish in color within a week (Fig. 4b) and hatched within two weeks (Fig. S2). Meanwhile, some field-collected eggs exhibited atypical black coloration (Fig. 4c), from which tiny black wasps emerged. The undescribed wasp species was named *Trissolcus brevinotaulus* based on the specimens we obtained (39) (Fig. 4d). When we marked and monitored 50 egg masses consisting of 223 eggs in the field, more than half of the eggs were parasitized by *T. brevinotaulus* (Fig. 4e), uncovering high levels of wasp parasitism in nature. In an attempt to evaluate the effect of egg-covering fungi on wasp parasitism, we generated fungus-removed egg masses by brushing and fungus-covered egg masses without brushing, and presented them to reproductive female wasps. In the experimental arena, the female wasps approached to both the fungus-removed eggs and the fungus-covered eggs, but their behaviors were markedly different. On the fungus-removed eggs, the female wasps exhibited oviposition behavior immediately after careful antennal drumming on the egg surface (Movie S3). On the fully fungus-covered eggs, by contrast, the female wasps were unable to approach to the eggs for oviposition (Movie S4). On the sparsely fungus-covered eggs, notably, the female wasps pressed down the hyphae by active antennal drumming and finally, though not always, oviposited successfully (Movie S5). Note that antennae of *T. brevinotaulus* are conspicuously thicker in females than in males, which are plausibly not only for sensing host eggs but also for flattening the hyphal thicket (Fig. S10a-d). Concordantly, the fungus-removed eggs suffered significantly higher wasp parasitism than the fungus-covered eggs (Fig. 4f). In addition, we experimentally generated fungus-suppressed egg masses by ablation of maternal hindlegs (Fig. S10e-h), and confirmed that the fungus-suppressed eggs suffered more oviposition trials and higher parasitism success than the fungus-covered eggs (Fig. 4g, h). It was impressive that the wasps did not avoid but actively approached and contacted the fungal hyphae, which entailed frequent self-grooming behaviors (Movies S3-S5). Even when the wasps were experimentally forced to contact with plenty of conidia of the egg-covering fungi, the survival of the fungus-treated wasps was not different from the survival of the untreated control wasps (Fig. 4i). These observations favor the idea that the nature of fungal defense is physical rather than chemical or pathogenic.

**Fig. 4.**
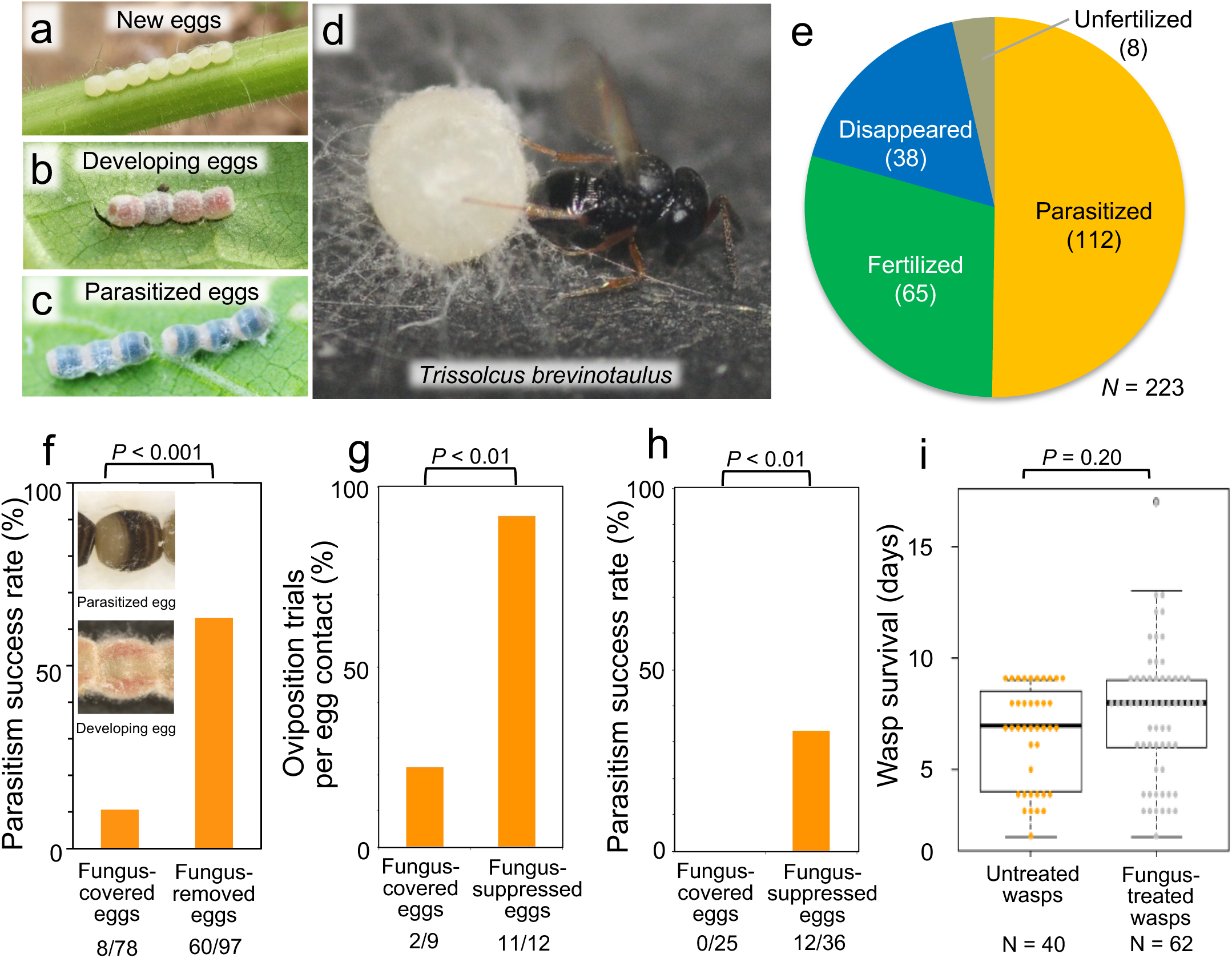
Suppression of wasp parasitism by egg-covering fungal hyphae. (**a**) Newly laid eggs white in color. (**b**) Developing fertilized eggs reddish in color. (**c**) Wasp-parasitized eggs black in color. (**d**) *T. brevinotaulus* ovipositing into an egg of *M. gracilicorne*. (**e**) Wasp parasitism rate in natural eggs of *M. gracilicorne*. In total, newly laid 50 egg masses consisting of 223 eggs were marked in the field, and inspected one week later. (**f**) Parasitism success rates of *T. brevinotaulus* on fungus-covered eggs and fungus-removed eggs by brushing. The fungus removing procedure is shown in Movie S3. (**g, h**) Oviposition trials per egg contact (**g**) and parasitism success rates (**h**) of *T. brevinotaulus* on fungus-covered eggs and fungus-suppressed eggs produced by ablation of maternal hindlegs. The fungus suppression procedure is shown in Fig. S10e-h. (**i**) Survival of *T. brevinotaulus* experimentally forced to contact with conidia of the hindleg-derived fungi. Statistical significance levels are estimated by Chi square test for (**f-h**) and generalized linear model test with a Poisson distribution for (**i**).

### Discovery of defensive fungal symbiosis mediated by specialized organ on insect hindleg

On the basis of these results, we conclude that the fungi selectively cultured on the female’s hindleg organ are transferred to eggs and play a defensive role against wasp parasitism, uncovering a previous unknown type of external symbiotic organ consisting of highly concentrated glandular mycangia for fostering the specific fungal associates. At least against the egg parasitoid *T. brevinotaulus*, the nature of defense is structural in that the egg-covering fungal hyphae physically distract approaching wasps and prevent parasitism. Notably, it was reported that the common cutworm moth *Spodoptera litura* lay eggs densely covered with scale hairs, which prevent egg parasitism by the wasp *Trichogramma chilonis* (40). Such structural defense against natural enemies is also known from gall-forming insects, in which thick and voluminous gall tissues, often with conspicuous projections or hairs on the surface, prevent access and oviposition of parasitoid wasps (41). In some gall midges, their symbiotic fungi contribute to the hardness of gall wall as well as serve as the nutritional source for larvae (42,43). Previous studies on defensive microbial symbioses in diverse insects have identified antibiotics (28,30–33), polyketide toxins (44,45), phage-encoded toxins (46), ribosome inactivating proteins (47) and other bioactive substances as symbiont-derived chemical factors for defense (36–38). Although not detected in this study, the possibility of chemical defense by the hindleg fungi against hitherto unidentified natural enemies of *M. gracilicorne* should be pursued in future studies.

### Evolutionary origin of defensive fungal symbiosis on specialized hindleg organ

The evolutionary origin of the hindleg organ and the defensive fungal symbiosis in *M. gracilicorne* remains to be an enigma. Enlarged hindlegs are known from several groups of heteropteran bugs (ex. Coreidae and Alydidae), but they are conspicuous in males and used for courtship fighting and territorial behaviors (48,49). Female-specific exaggerated hindlegs seem unique to the family Dinidoridae, and the conspicuous morphological trait is conserved across two major dinidorid lineages Dinidorinae (∼70 species) and Megymeninae (∼24 species including *M. gracilicorne*) (5–8). Hence, the hindleg organ is likely to have evolved in the common ancestor of the Dinidoridae. We hypothesize, although speculative, that the female-specific hindleg organ evolved originally for smearing some secretion onto eggs for chemical defense or camouflage, but environmental entomopathogenic fungi colonized the organ and exploited the secretion, and the insects established specific association with benign fungal species and co-opted them for egg defense. To test this hypothesis, the structure of the hindleg organs, associated microorganisms, and maternal behaviors upon oviposition should be comparatively investigated among diverse dinidorid stinkbugs in the world.

### Relevance to evolutionary theory of symbiont transmission mode

In *M. gracilicorne*, the specific fungal associates are vertically transmitted to eggs by female’s specialized structures and behaviors, but the association is cancelled during the nymphal development and restarted by newly-emerged adult females. At a glance, it was enigmatic why elaborate vertical symbiont transmission is combined with environmental symbiont acquisition in the insect life cycle. Our discovery of the symbiont’s defensive role against the egg parasitoid resolved the enigma – the maternal fungal smearing onto eggs is actually not for transmission but for defense. In principle, vertical symbiont transmission through host generations favors host-symbiont cooperation and facilitates the evolution of mutualism, whereas horizontal symbiont acquisition generally facilitates symbiont’s virulence and counters the evolution of mutualism (50–52). Meanwhile, theoretical studies predicted that mutualism without vertical transmission may evolve under the following conditions: (i) vertical transmission of the symbiont incurs some cost for the host, (ii) exploitation by the symbiont negatively affect the host, (iii) the host controls the vertical transmission process, and (iv) the host is able to discriminate benevolent symbionts from parasitic ones (53–55). Notably, all these conditions seem to apply to the fungal symbiosis in *M. gracilicorne*.

### Impact on transmission mode of coexisting symbiont

Here we point out that the external fungal symbiont may have evolutionarily impacted on a coexisting microbial symbiont in *M. gracilicorne*. In general, stinkbugs of the superfamily Pentatomoidea are dependent on specific γ-proteobacterial symbionts localized to the posterior midgut, and the beneficial/essential symbionts are vertically transmitted from mother to offspring via egg surface smearing, symbiont-containing capsules, symbiont-containing gelatinous secretion, or other means (13–15) (Fig. S11). However, our recent study uncovered that the beneficial gut symbiont of *M. gracilicorne* is not transmitted vertically but acquired from the environment every generation (56), which is exceptional among the Pentatomoidea (Fig. S11). It seems plausible, although speculative, that the egg-covering defensive fungi disturbed vertical transmission of the gut symbiotic bacteria and have promoted the evolution of environmental symbiont acquisition in an ancestor of the Dinidoridae.

### Conclusion and perspective

Our revisiting of the female-specific “tympanal organ”, which had been known from dinidorid stinkbugs for decades, led to the unexpected discovery of novel external fungal symbiosis for physical defense against wasp parasitism. The defensive fungal symbiosis on insect hindlegs provides an impressive case as to how evolutionarily novel traits for microbial symbiosis emerge, develop, and constitute the elaborate syndrome that integrates molecular, cellular, morphological and behavioral specializations. The bizarre coexistence of vertical symbiont transmission, symbiont loss, and environmental symbiont re-acquisition in the insect life cycle can be accounted for by benefit of the fungal egg defense and potential cost of nymphal infection, which highlights theoretical and empirical relevance of symbiont transmission mode to parasitism-mutualism evolutionary continuum. Future studies should focus on the molecular mechanisms underpinning the development of the peculiar symbiotic organ and the specificity of the fungal cultivation on the symbiotic organ, for which transcriptomic, metabolomic and RNAi screening approaches on morphogenesis and functioning of the hindleg organ will provide important clues.

## Materials and Methods

Detailed Materials and Methods are available in Supplementary Information.

## Acknowledgements

We thank João Araújo and Hassan Salem for reading and constructively commenting on the manuscript. This study was supported by the Japan Society for the Promotion of Science (JSPS) KAKENHI Grant JP17H06388 to TF and RK, and the Japan Science and Technology Agency (JST) ERATO Grant JPMJER1902 to TF. TN was supported by a JSPS Research Fellowship for Young Scientists.

## Author contributions

TH and TF conceived the study. TN, HM and MM conducted the majority of experimental works, where TN mainly conducted microbiological, histological and field experiments, HM mainly performed behavioral experiments, and MM mainly carried out transcriptomic, proteomic and biochemical experiments. TN, TH and MT isolated and analyzed the diverse fungal strains. ST discovered the peculiar behavior of *M. gracilicorne*. NN, RK and MM constructed molecular phylogeny of the host stinkbugs. TF wrote the paper with input from all authors.

## Supplementary Information

### Materials and Methods

#### Insect collection and rearing

The samples of *M. gracilicorne* subjected to fungal isolation and characterization, fungal ITS cloning and sequencing, and fungal ITS amplicon sequencing are listed (Table S1). For other experimental purposes, the insects were mainly collected at Chikusei, Ibaraki, Japan from *Sicyos angulatus*, and used immediately for experiments, or maintained on fresh cucumber fruits and distilled water at 25°C under a 16 h light and 8 h dark condition as described previously (56).

#### Fungal isolation and characterization

Each field-collected adult female was individually reared in a plastic container, where the first egg mass and the second egg mass were sampled. Then, the insect was subjected to sampling of the hindlegs, from which fungal isolation was conducted. Each egg mass was individually kept in a humidified petri dish for a week, which were massively covered with fungal hyphae, and was subjected to fungal isolation. Each sample was scraped with a sterilized platinum wire, spread on PDA agar plates, and cultured at 25°C under constant darkness. After several days of incubation, four colonies were randomly picked and spread on fresh PDA agar plates, by which up to four fungal isolates per sample were purified and established. The fungal isolates were cultured on coverslips, mounted in lactophenol, and morphologically observed under a phase contrast microscope (Axiophot, Zeiss).

#### Fungal ITS sequencing and analysis

The fungal isolates were subjected to DNA extraction using a liquid nitrogen freeze-crushing mill (SK-200, Tokken) and QIAamp DNA Mini Kit (QIAGEN), PCR amplification of fungal ITS region with the primers ITS4 (5‘-TCC TCC GCT TAT TGA TAT GC-3’) and ITS5 (5’-GGA AGT AAA AGT CGT AAC AAG G-3’), and sequencing of the PCR products with the same primers using BigDye Terminater v 3.1 Cycle Sequencing Kit and ABI PRISM 373 DNA Sequencer (Applied Biosystems) or outsourced (Eurofin Genomics). The fungal ITS sequences were subjected to BLAST searches and molecular phylogenetic analyses. Multiple alignments of the nucleotide sequences were generated using MAFFT (57), from which ambiguously aligned sites and gap-containing sites were removed using trimAl v1.2 (58). Maximum-likelihood phylogenies were constructed using RAxML-ng v1.2.1 (59) with 1,000 bootstrap resamplings, while Bayesian phylogenetic inference was conducted using MrBayes v3.2.7a (60). Substitution model selection was conducted under the Akaike information criterion using modeltest-ng v0.1.7 (61). The nucleotide sequence data obtained in this study were deposited in the DNA Data Bank of Japan (DDBJ) under accession numbers LC792825 – LC793396 (Table S1).

#### Fungal ITS amplicon sequencing and analysis

After fungal isolation, each of the hindleg samples and the egg samples was subjected to DNA extraction as described above, and PCR amplification of the fungal ITS2 region with the primers ITS3 (5’-GCA TCG ATG AAG AAC GCA GC-3’) and ITS4 using Phusion High-Fidelity PCR Master Mix (New England Biolabs). The PCR products were purified by Qiagen Gel Extraction Kit (Qiagen) and subjected to library construction using NEBNext UltraTM DNA Library Prep Kit for Illumina (New England Biolabs). The 250-bp paired-end sequencing of the libraries using HiSeq 2500 or NovaSeq 6000 (Illunmina) was outsourced (Filgen). The adaptor, primer, and low-quality sequences were removed using Cutadapt v2.8 (62). Subsequently, the trimmed reads were analyzed on the QIIME2 platform (63), in which taxonomic classification was conducted based on the UNITE eukaryote ITS database v9.0 (64). Sequence variants with more than 99% similarity were defined as an operational taxonomic unit (OUT) through the DECIPHER package (65), and subjected to the statistical processing using R v4.3.2 (66). The raw sequence data were deposited in the DDBJ Sequence Read Archive under accession numbers DRX509842 – DRX509916 (Table S1).

#### Histological observation of hindleg organ

Hindlegs were excised from field-collected mature females and males, and also from laboratory-reared immature females within seven days after eclosion. For inspection of the external fine structures, each hindleg sample was fixed on the observation stage with a double-coated adhesive tape, gold-coated using a vapor deposition equipment (Smart Coater, JEOL), and observed under a scanning electron microscope (JCM-6000, JEOL). For observation of the tissue sections, the excised hindlegs were fixed in 10% (v/v) formalin in ethanol for 1 h, dehydrated with ethanol, embedded in a methacrylate resin (Technovit 7100, Kulzer), sectioned at 4 μm thickness on a microtome (RM2165, Leica), adhered on glass slides, subjected to periodic acid-Schiff (PAS) staining, hematoxylin-eosin staining and/or toluidine blue staining, mounted in Entellan New (Sigma-Aldrich), and observed under a light microscope (Axiophot, Zeiss). For observation of the fine structures of the hindleg organ, the dissected hind tibiae were prefixed with 2.5% glutaraldehyde in 0.1 M phosphate buffer (pH 7.4) at 4°C overnight, and postfixed with 2% osmium tetroxide in 0.1 M phosphate buffer (pH 7.4) at 4°C for 1 h, dehydrated through a water-ethanol series, embedded in Epon812 resin, sectioned on an ultra-microtome (EM UC7, Leica), mounted on copper meshes, stained with uranyl acetate and lead citrate, and observed under a transmission electron microscope (H-7600, Hitachi).

#### Transcriptomic analysis of hindleg organ

Midlegs and hindlegs were sampled from laboratory-reared mature males, laboratory-reared mature females whose hindleg organs were fully covered with fungal hyphae, and also from laboratory-reared immature females within seven days after eclosion whose hindleg organs exhibited no visible fungal growth. The leg samples were subjected to RNA extraction using a liquid nitrogen freeze-crushing mill (SK-200, Tokken) and RNAiso (Takara) and chloroform. The RNA-containing supernatants were purified by RNeasy Mini Kit (Qiagen) and subjected to cDNA library construction using NEBNext UltraTM DNA Library Prep Kit for Illumina (New England Biolabs). Sequencing of the libraries using HiSeq 2500 (Illumina) was outsourced (Dojin Chemical). The RNA sequencing data were analyzed through the following pipeline. After low-quality read trimming by Trimmomatic v3.6 (67), sequence data were subjected to *de novo* assembling using Trinity v2.11.0 (68). Read mapping and expression level estimation were performed by Salmon v1.5.1 (69). Because the obtained reads were derived from the insects, fungi, and a small amount of contaminated bacteria, we determine the contig origins based on similarity scores of BLAST search against several protein databases; we classified contigs with the highest score against RefSeq Hemiptera or RefSeq *Drosophila melanogaster* proteins as insect-derived genes, those for Swiss-Prot fungal proteins as fungal genes, and those for Swiss-Prot bacterial proteins as bacterial genes. We annotated gene functions with the aid of Trinotate program (70) in addition to the BLAST results. Differentially expressed genes between tissues were statistically determined by edgeR algorism (71) on R v4.3.2 (66).

Functional enrichment analysis was performed using DAVID platform (72). The raw RNA sequencing data were deposited in the DDBJ Sequence Read Archive under accession numbers DRX509810 – DRX509841 (Table S4).

#### Histological observation and proteomics of egg surface

Surface structures of newly laid white eggs (0-day-old) and developing reddish eggs (7-day-old) were observed by scanning electron microscopy as described above. Eggshells and also mature oocytes in the ovaries were subjected to tissue sectioning and PAS staining for visualizing the polysaccharide-rich surface layer as described above. A variety of solvents were attempted to dissolve the surface layer (see Fig. S8), of which SDS-PAGE sample buffer (50 mM Tris-HCl, 2% SDS, 0.6% 2-mercaptoethanol, pH 6.8) turned out to be the best. The extracted samples were treated with methanol-chloroform-water mixture to remove SDS and to precipitate proteins. After dried in vacuo, the precipitates were subjected to a phase transfer surfactant-aided trypsin digestion (73). The samples were resuspended to 100 mM Tris-HCl buffer (pH 9.0) containing 12 mM sodium deoxycholate and 12 mM sodium lauroylsarcosinate, and heated at 95°C for 5 min. Reduction and alkylation of cysteine residues were performed by reacting with 20 mM dithiothreitol and 40 mM iodoacetamide, respectively. Then, the samples were subjected to protein digestion using Trypsin/Lys-C Mix (Promega) for 16 h at 37°C. After removal of the surfactants by liquid-phase extraction using ethyl acetate, the digested peptides were purified with a GL-Tip SDB (GL Science) column. The peptide mass analysis was carried out with an LC-MS system composed of an ultrahigh performance liquid chromatograph (ACQUITY UPLC H-Class, Waters) and a high-resolution mass spectrometer (Xevo G2-XS QToF, Waters) connected to an electrospray ionization source. Peptides were separated on a CSH column (2 mm i.d., 150 mm, Waters), and their precursor and fragment mass data were obtained using a positive MS^E^ mode. Protein identification was performed on ProteinLynx Global Server (Waters) by reference to the protein sequences deduced from the transcriptome analysis.

#### Field observation and collection of egg parasitic wasp

From the end of June to July in 2014, we performed regular field monitoring, inspection and sampling of egg masses of *M. gracilicorne* at Chikusei, Ibaraki, Japan. Every week we visited a bush of *C. angulatus* on the riverbank of Kokai River, where *M. gracilicorne* was found at an exceptionally high density, to monitor natural egg masses laid on the plant vines and leaves. Young white eggs (see Fig. 4a) were marked and recorded, and inspected and collected next week. We checked 50 egg masses consisting of 223 eggs in total, and judged each egg condition as developing (turning reddish, see Fig. 4b), parasitized (turning blackish, see Fig. 4c), unfertilized (remaining white), or disappeared. Reddish developing eggs and blackish parasitized eggs were collected immediately and kept in humidified petri dishes in the laboratory, from which emerging parasitic wasps, *T. brevinotaulus*, were collected. The adult wasps were placed in small plastic cups (40 mm in diameter, 25 mm high) with filter paper on the bottom and fed with 50% honey water. The wasps were maintained at 25°C under a long-day regime of 16 h light and 8 h dark until used for experiments.

#### Preparation of fungus-covered eggs and fungus-removed eggs for wasp parasitism assay

Egg masses laid on filter paper were collected from a group of mass-reared adult insects of *M. gracilicorne*, and were individually transferred to and kept in humidified plastic cases (40 x 40 mm, 15 mm high). For evaluating the degree of fungal growth, each plastic case was lined at the bottom with moistened black filter paper. The eggs were maintained at 25°C under a long-day regime of 16 h light and 8 h dark for three days. Of the fungus-covered egg masses, 14 egg masses (consisting of 3-8 eggs) were used as control fungus-covered egg masses, whereas 19 egg masses (consisting of 3-7 eggs) were treated as fungus-removed egg masses by brushing and removing the fungal hyphae (see Movie S3). The experimental egg masses were subjected to wasp parasitism assay. In each petri dish arena, an experimental egg mass was presented to a female wasp for 1 h. These eggs were maintained and diagnosed as parasitized or not based on egg color (see Fig. 4f) ten days after the treatment.

#### Preparation of fungus-covered eggs and fungus-suppressed eggs for wasp parasitism assay

Field-collected reproductive females of *M. gracilicorne* were randomly allocated to the following experimental treatments, namely no treatment control, midlegs-amputated treatment, and hindlegs-amputated treatment (Fig. S10e-g). Each female was paired and maintained with a male in a plastic petri dish (90 mm in diameter, 25 mm high) at 25°C under a long-day regime of 16 h light and 8 h dark for three days. Then, the females were individually reared and allowed to lay egg masses, by which 35 egg masses were obtained from 11 females in the control group, 32 egg masses from 13 females in the midlegs-amputated group, and 33 egg masses from 12 females in the hindlegs-amputated group. In the control group and the midlegs-amputated group, most of the egg masses were covered with fungal hyphae. In the hindlegs amputated group, by contrast, more than half of the egg masses grew little fungal hyphae (Fig. S10e-h). Four fungus-covered egg masses (consisting of 6-11 eggs) from the midlegs-amputated group and four fungus-suppressed egg masses (consisting of 5-8 eggs) from the hindlegs-amputated group were subjected to the wasp parasitism assay around a week after oviposition. In each petri dish arena, an experimental egg mass was presented to five female wasps and continuously observed for 1 h, during which wasps approaching to, climbing on, and trying to oviposit into the eggs were confirmed. We observed that the wasps approached to 9 fungus-covered eggs and 11 fungus-suppressed eggs, respectively. Of these, the eggs to which the wasps tried to insert their ovipositor were judged as oviposited. The eggs were maintained and diagnosed as parasitized based on wasp emergence.

#### Pathogenicity assay of egg-covering fungi against parasitic wasps

Hyphae covering an egg mass was inoculated to and cultured on PDA agar plates at room temperature for 10 days, on which the fungi grew fully and formed plenty of conidia. In total 62 adult wasps of *T. brevinotaulus* were introduced into the inoculated agar plates, incubated for 1 h, transferred to small clean petri dishes, and individually maintained with 50% honey water as food until they died. For control, 40 adult wasps were incubated in sterile PDA agar plates for 1 h, and maintained and inspected in the same way.

#### Mitochondrial genome sequencing and phylogenetic analysis

The genomic DNA of *M. gracilicorne* was extracted with QIAamp DNA Mini Kit (QIAGEN). The DNAseq library was constructed by using Ligation Sequencing Kit (Oxford Nanopore Technologies, UK) and the resultant library was sequenced on MinION R9.4.1 (Oxford Nanopore Technologies). The reads obtained was de novo assembled by using Minimap2 Ver. 2.23-r1111 (74), Miniasm Ver. 0.3-r179 (75) and Minipolish Ver. 0.1.3 (76). The resultant contigs were polished with Medaka Ver. 1.0.3 (Oxford Nanopore Technologies). The contig corresponding to the mitochondrial genome of *M. gracilicorne* was identified based on the similarity to the *Nezara viridula* mitochondrial genome (NC_011755). The machine annotation of the mitochondria genome sequence was conducted with MITOS WebServer (http://mitos.bioinf.uni-leipzig.de/index.py) (77). The mitochondrial genome sequence of *M. gracilicorne* was subjected to molecular phylogenetic analyses together with 23 mitochondrial genome sequences representing 8 Pentatomoidea families with three outgroup taxa. The sequences were aligned using MAFFT v7.464 with L-INS-I algorithm (55) and ambiguously aligned sites were trimmed by using Gblocks 0.91b with manual inspection (78). A resultant concatenated alignment consisting of 3,539 amino acid sites representing 13 protein-coding genes was subjected to maximum likelihood (ML) analysis and Bayesian inference (BI) analysis. The optimal partitioning schemes and corresponding amino acid substitution models were estimated using PartitionFinder v2.1.1 (79). The ML analysis was conducted using IQ-TREE v2.0.3 (80). Bootstrap values were obtained with 1,000 resamplings. The BI analysis was conducted using MrBayes v3.2.7a (60) with 1,000,000 generations of Markov chain Monte Carlo runs with sampling every 100 generations. The first 25 % samples were discarded as burn-in and the remaining trees were used to calculate posterior probabilities. Stationarity was considered to be reached when the average standard deviation of split frequencies was below 0.01.

**Fig. S1.**
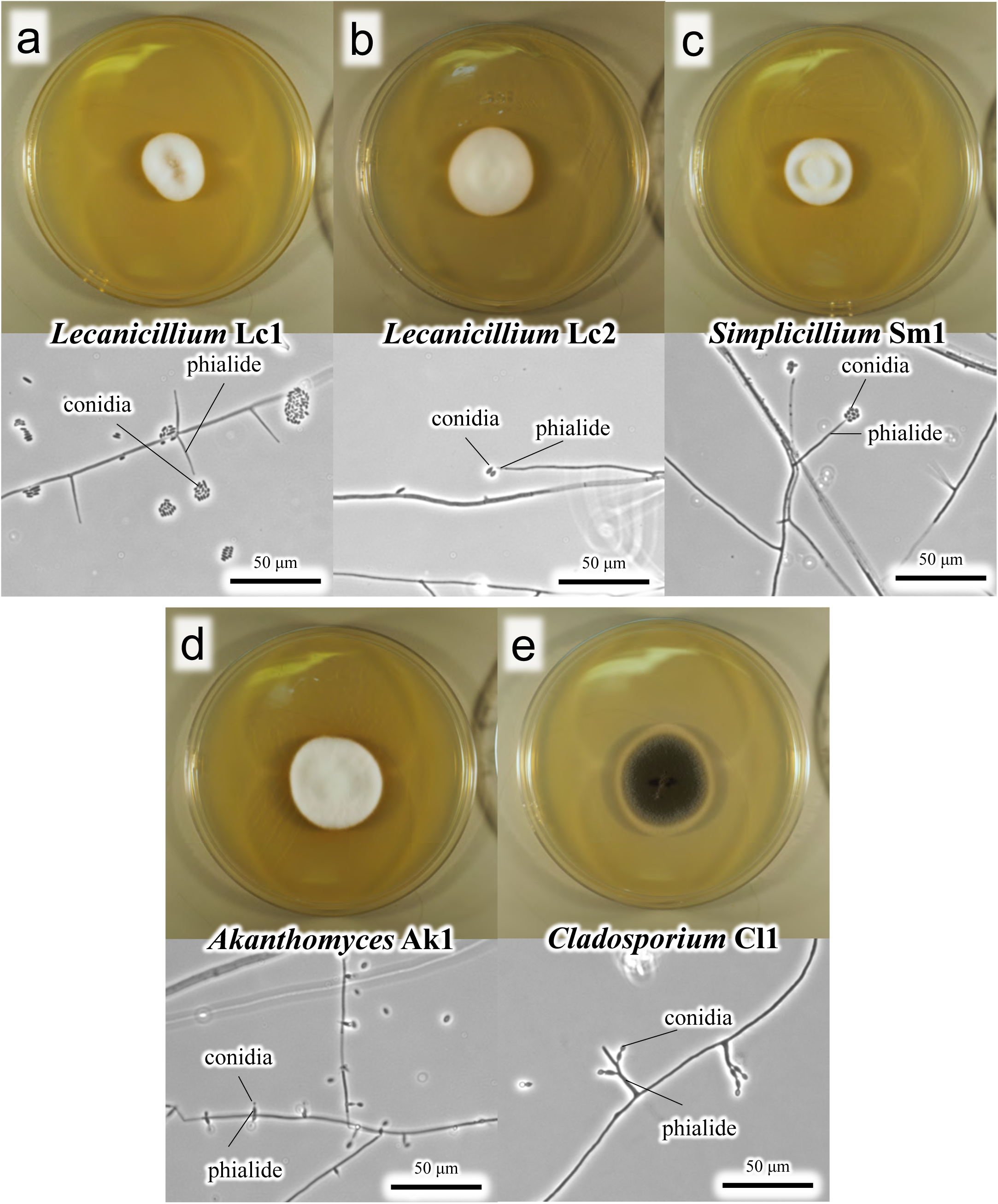
Fungi isolated from female’s hindleg organs and eggs of *M. gracilicorne*. (**a**) *Lecanicillium* Lc1. (**b**) *Lecanicillium* Lc2. (**c**) *Simplicillium* Sm1. (**d**) *Akanthomyces* Ak1. (**e**) *Cladosporium* Cla1. Note that (**a**)-(**d**) are frequently isolated, belonging to the same entomopathogenic group Cordycipitaceae, and regarded as dominant fungal associates, whereas (**f**) is isolated only sporadically and likely a fungal contaminant.

**Fig. S2.**
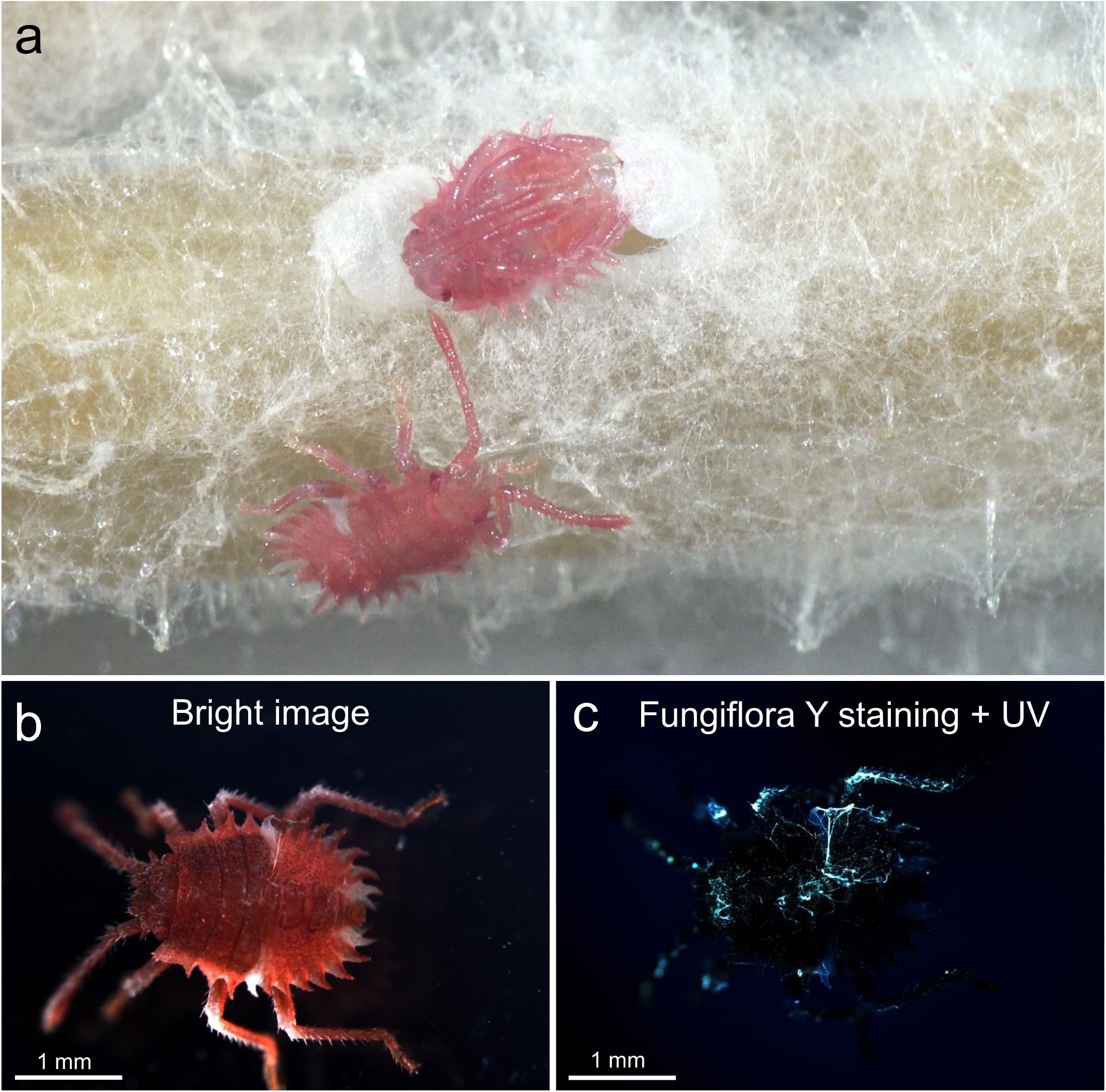
Newborn nymphs of *M. gracilicorne*. (**a**) Hatching nymphs. Also see Movie S2. (**b, c**) A newborn nymph under white light (**b**) and UV light (**c**) on which attaching fungal hyphae are stained with Fungiflora Y.

**Fig. S3.**
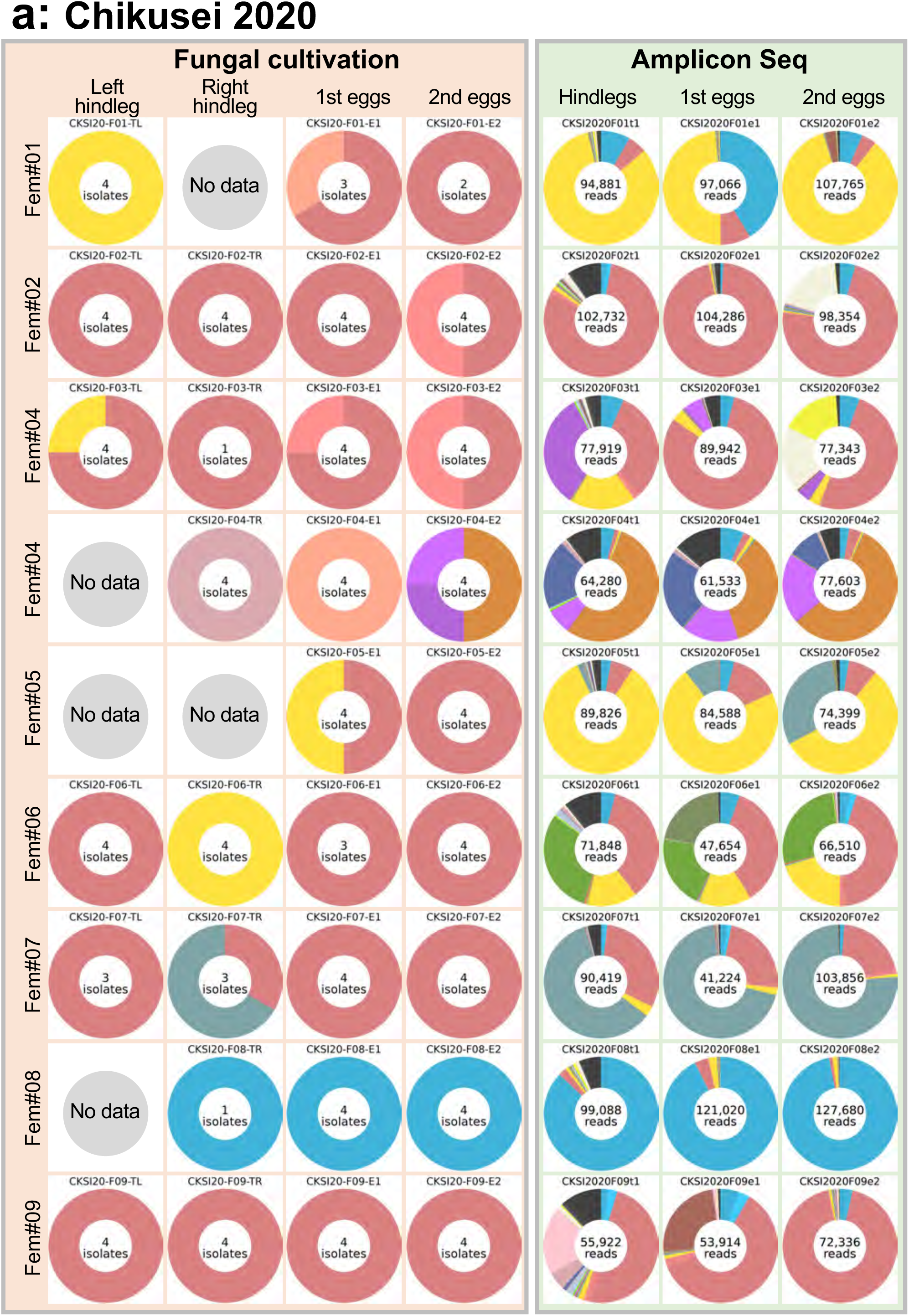
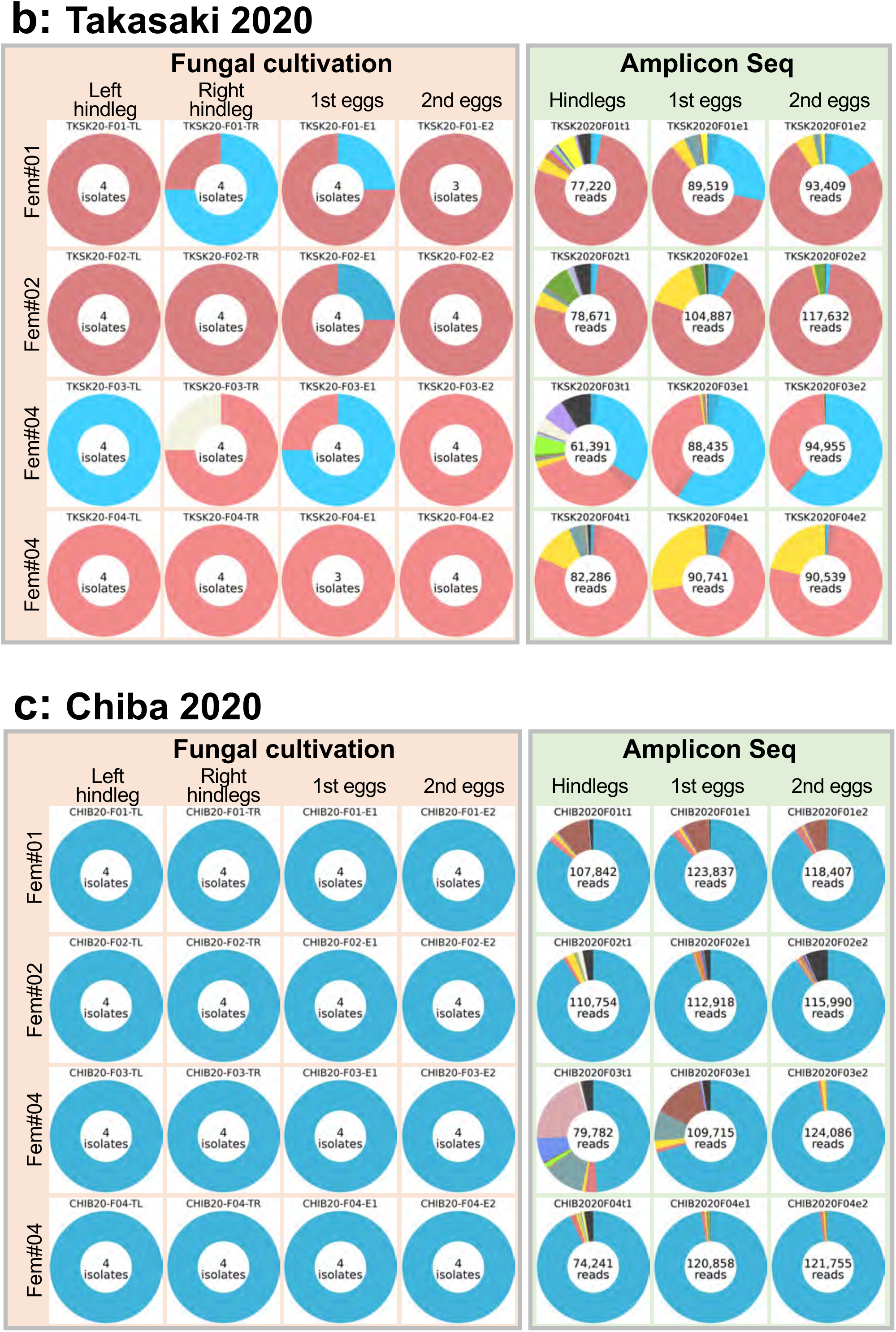
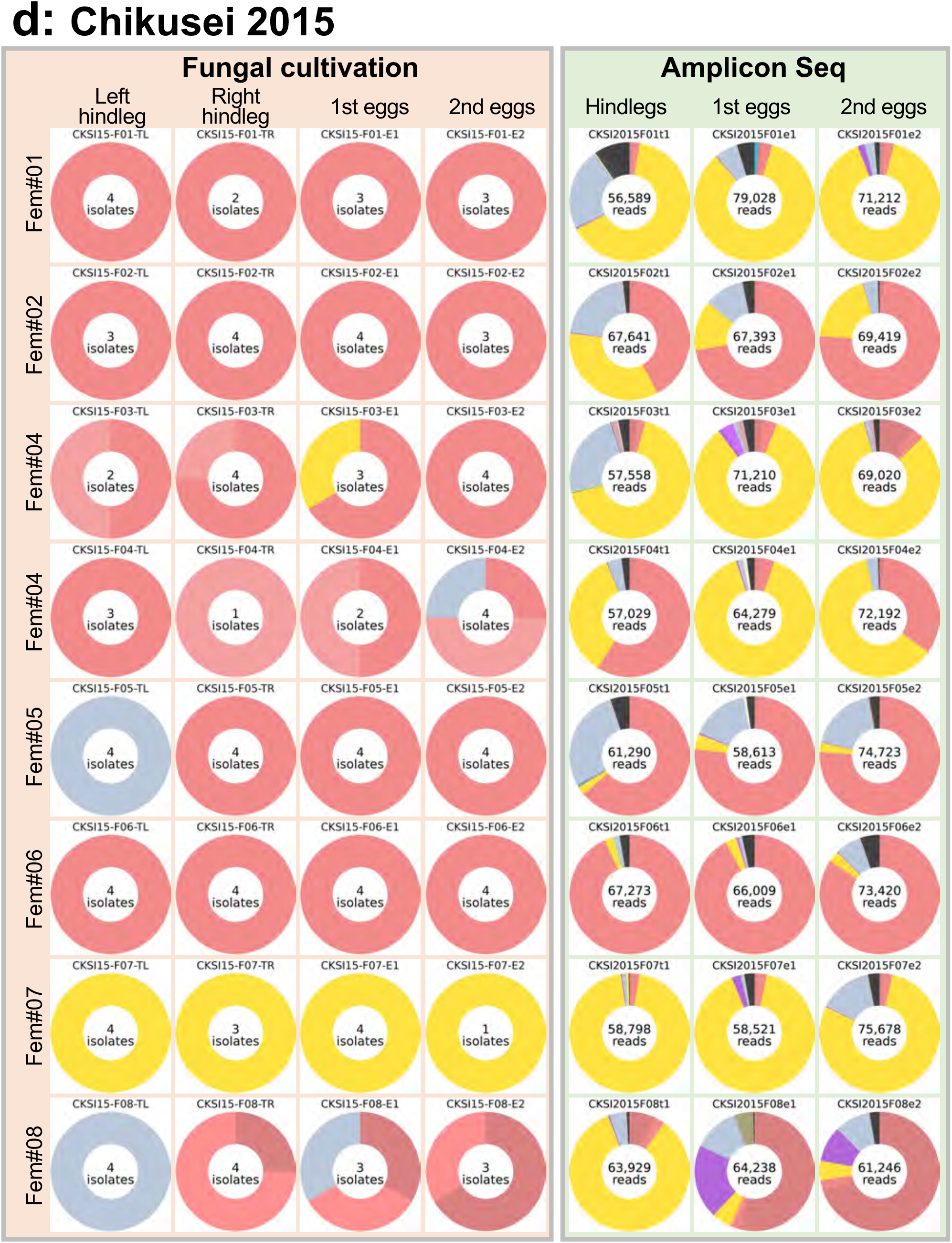
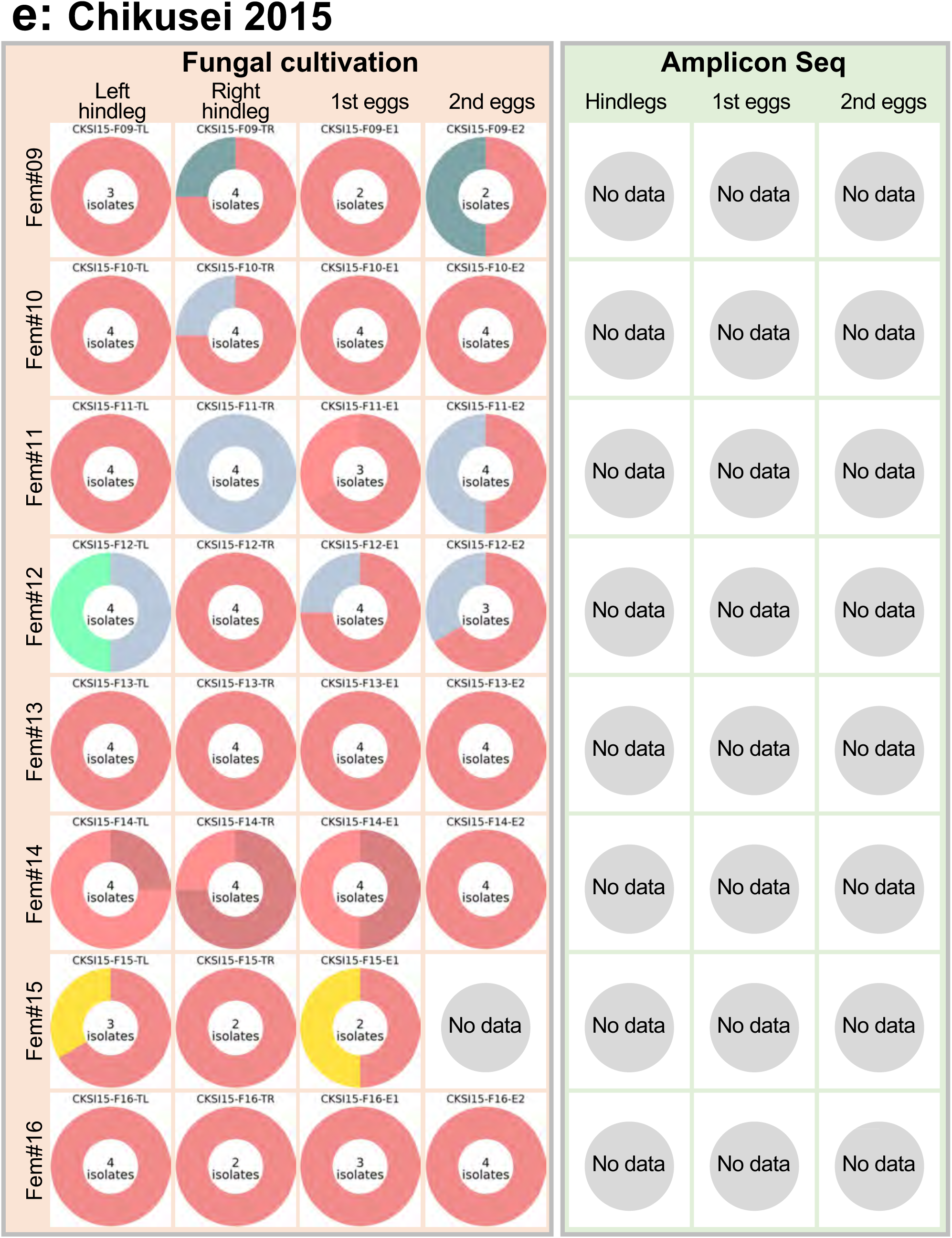
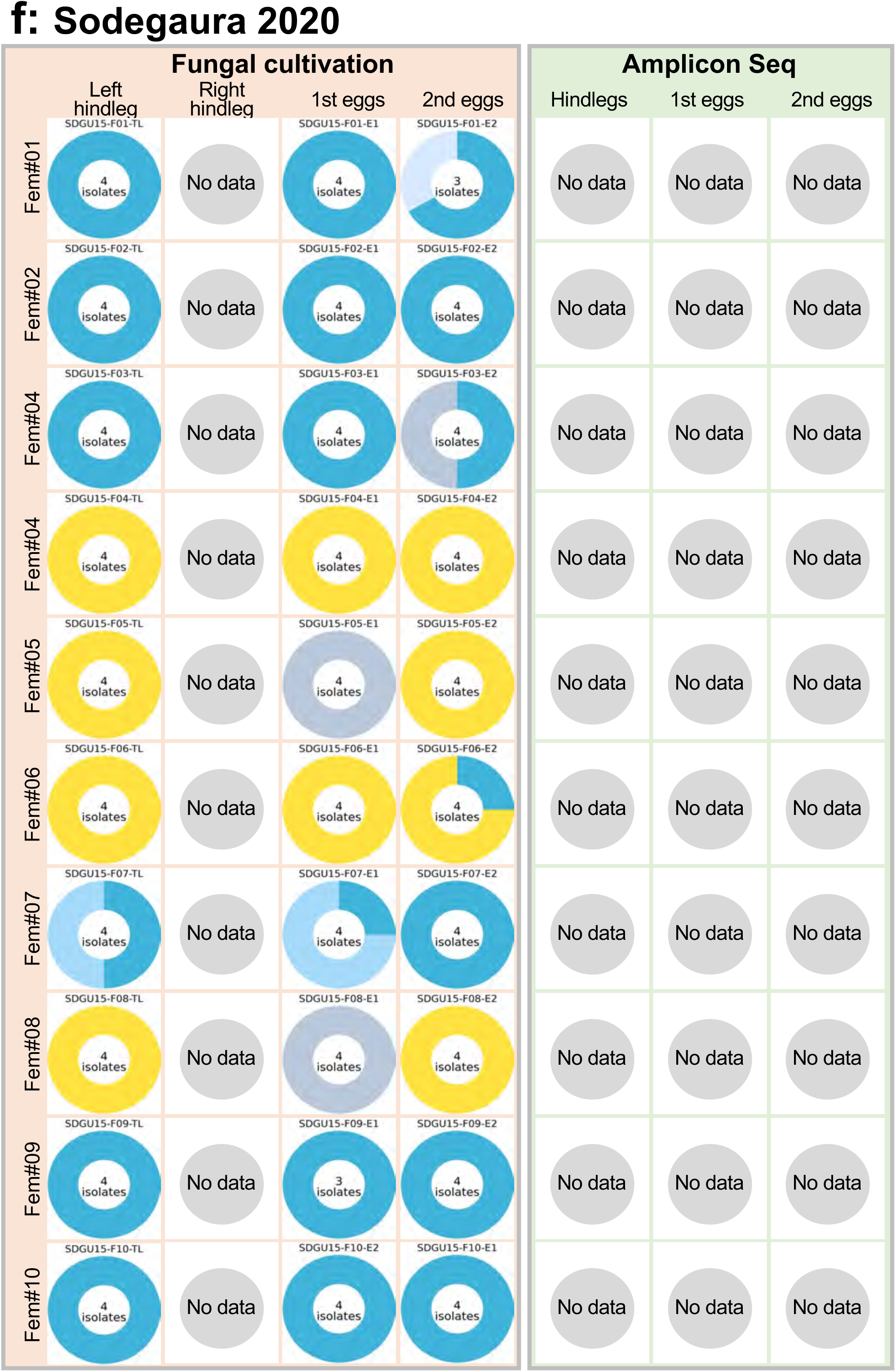
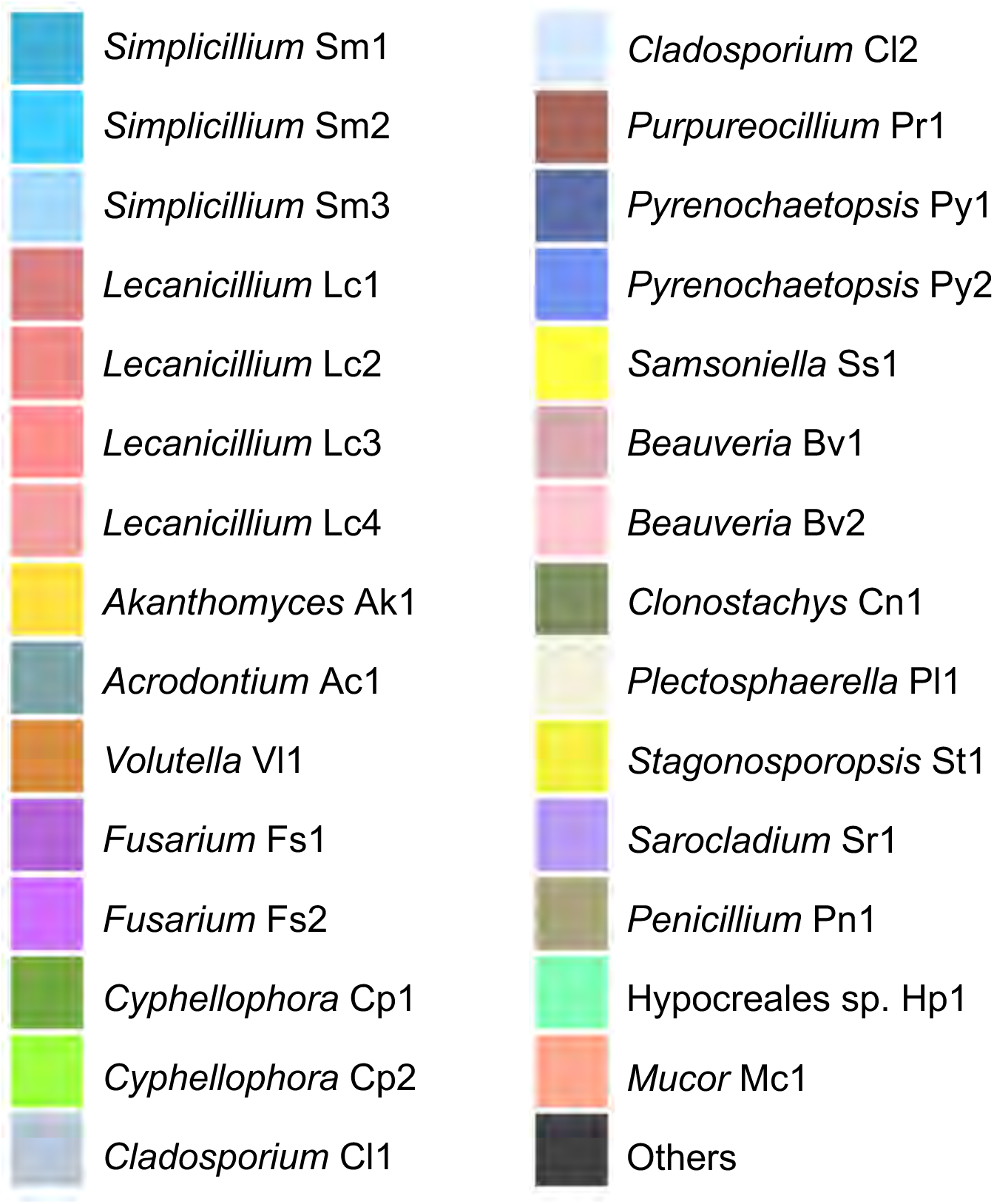
Fungal microbiota on female’s hindleg organs and eggs of *M. gracilicorne*. (**a**) Nine adult females collected at Chikusei, Ibaraki, in 2020. Eight adult females collected at Chikusei, Ibaraki, in 2015. (**b**) Four adult females collected at Takasaki, Gumma, in 2020. (**c**) Four adult females collected at Chiba, Chiba, in 2020. (**d**) Eight adult females collected at Chikusei, Ibaraki, in 2015. (**e**) Eight adult females collected at Chikusei, Ibaraki, in 2015. (**f**) Ten adult females collected at Sodegaura, Chiba, in 2015. For each insect, dissected hindlegs, first eggs and second eggs were subjected to fungal isolation, fungal DNA extraction, and PCR amplification and sequencing of ITS (left) and also direct DNA extraction and ITS amplicon sequencing (right) for (**a-d**), or fungal cultivation and ITS sequencing only for (**e, f**). Colored pie graphs depict major fungal strains detected. At the center of each pie graph, either the number of analyzed fungal isolates or the number of ITS amplicon sequencing reads are shown. For the localities, see the map in Fig. 2.

**Fig. S4.**
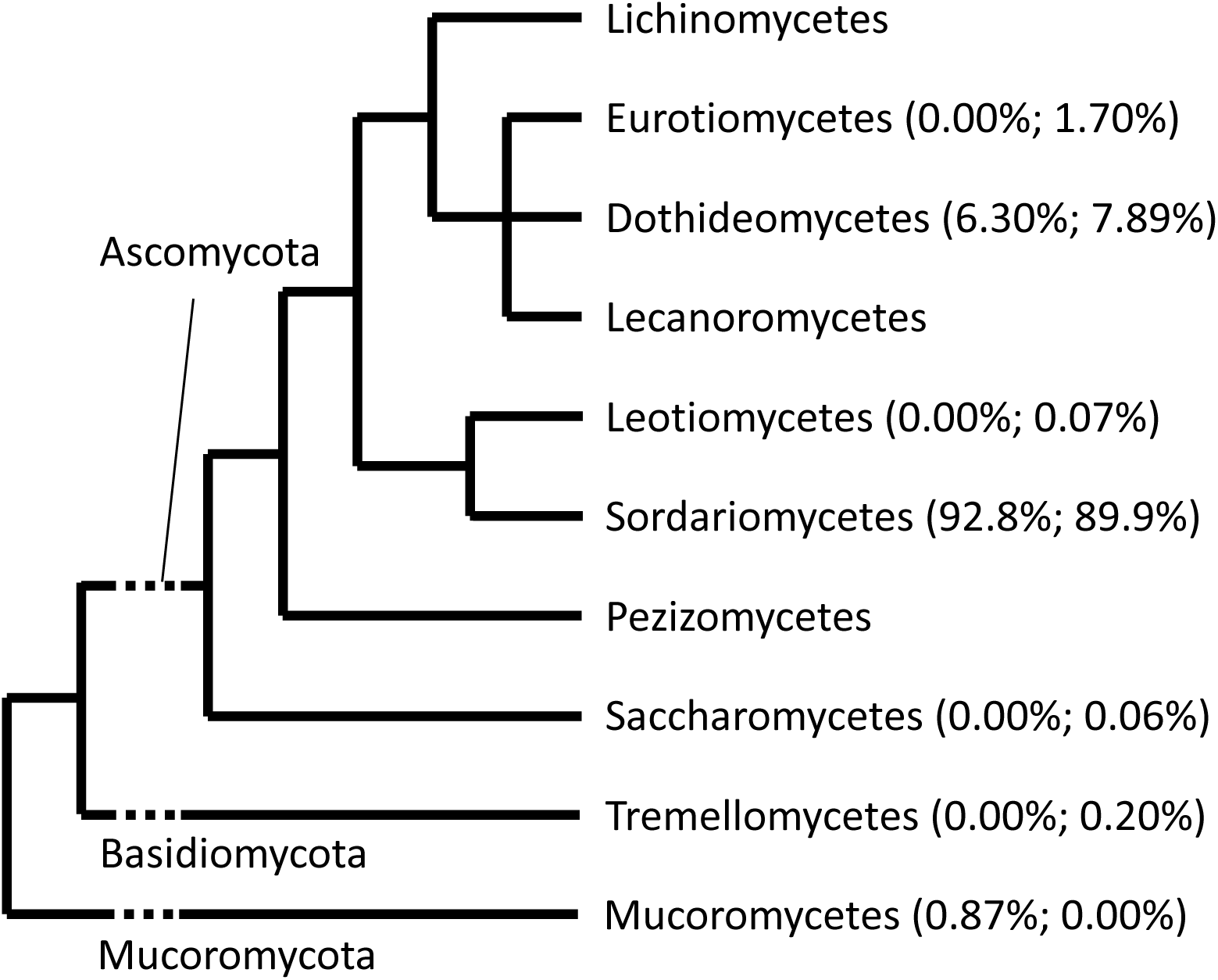
Phylogenetic placement of fungal groups detected from hindlegs and eggs of *M. gracilicorne*. Of 631 fungal isolates, fungal ITS sequences were successfully obtained for 572 isolates, of which 531 (92.8%) belong to the class Sordariomycetes whereas 36 (6.30%) are placed in the Dothideomycetes. Of 6,230,166 ITS seq reads obtained in total, 5,602,848 reads (89.9%) belong to the Sordariomycetes whereas 492,052 reads (7.89%) and 105,970 reads (1.70%) are placed in the Dothideomycetes and the Eurotiomycetes, respectively.

**Fig. S5.**
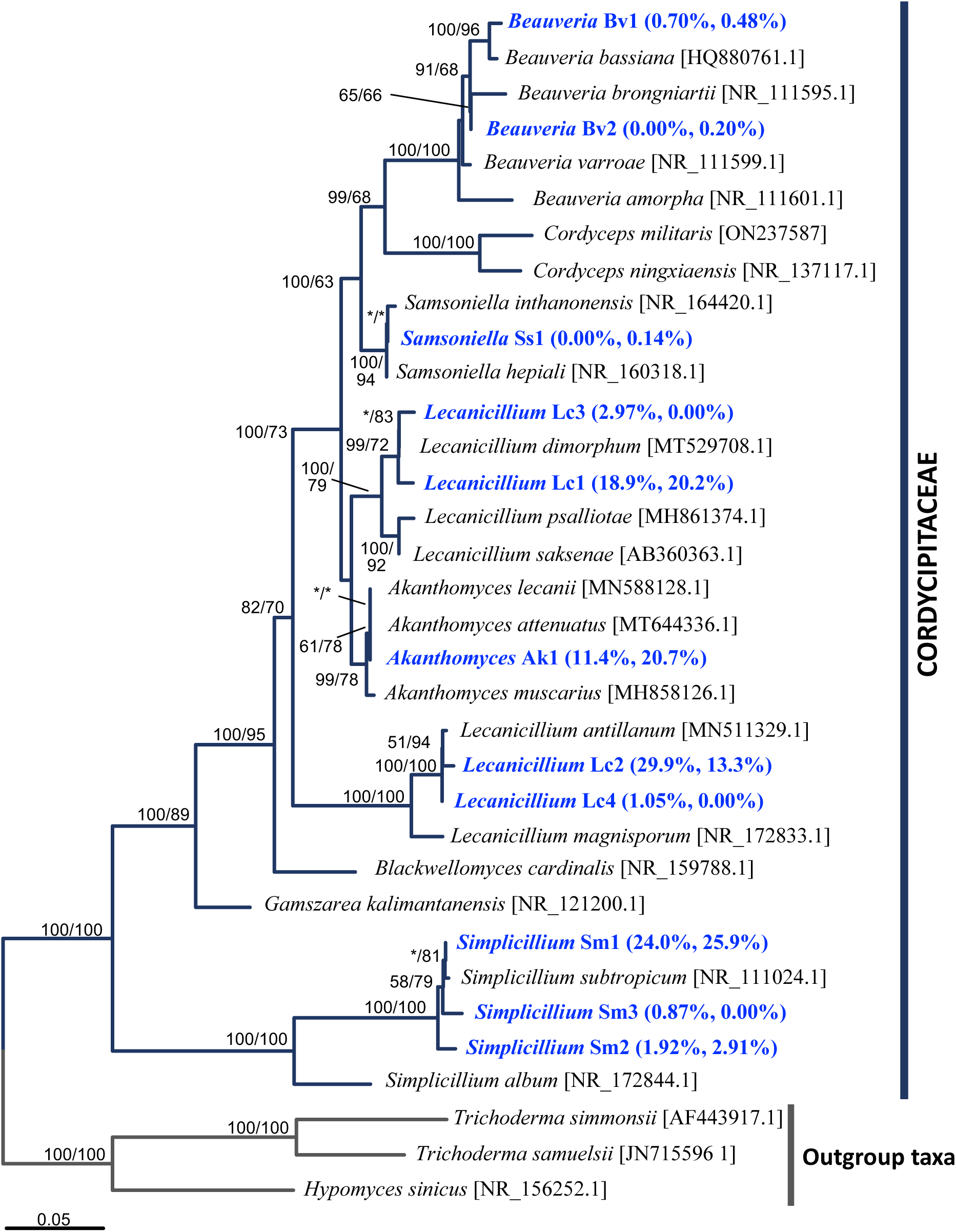
Phylogenetic placement of fungal strains detected from hindlegs and eggs of *M. gracilicorne* on the basis of fungal ITS sequences. A maximum likelihood phylogeny (ML) inferred from a total of 546 aligned nucleotide sites is shown, whereas Bayesian phylogeny (BY) exhibited substantially the same pattern. Statistical support values are shown at the nodes as bootstrap probabilities for ML/posterior probabilities for BY. Sequences obtained in this study were shown in blue, and each percentage of 572 isolates for fungal cultivation and 6,230,166 reads for ITS amplicon seq was indicated in parenthesis on the right side.

**Fig. S6.**
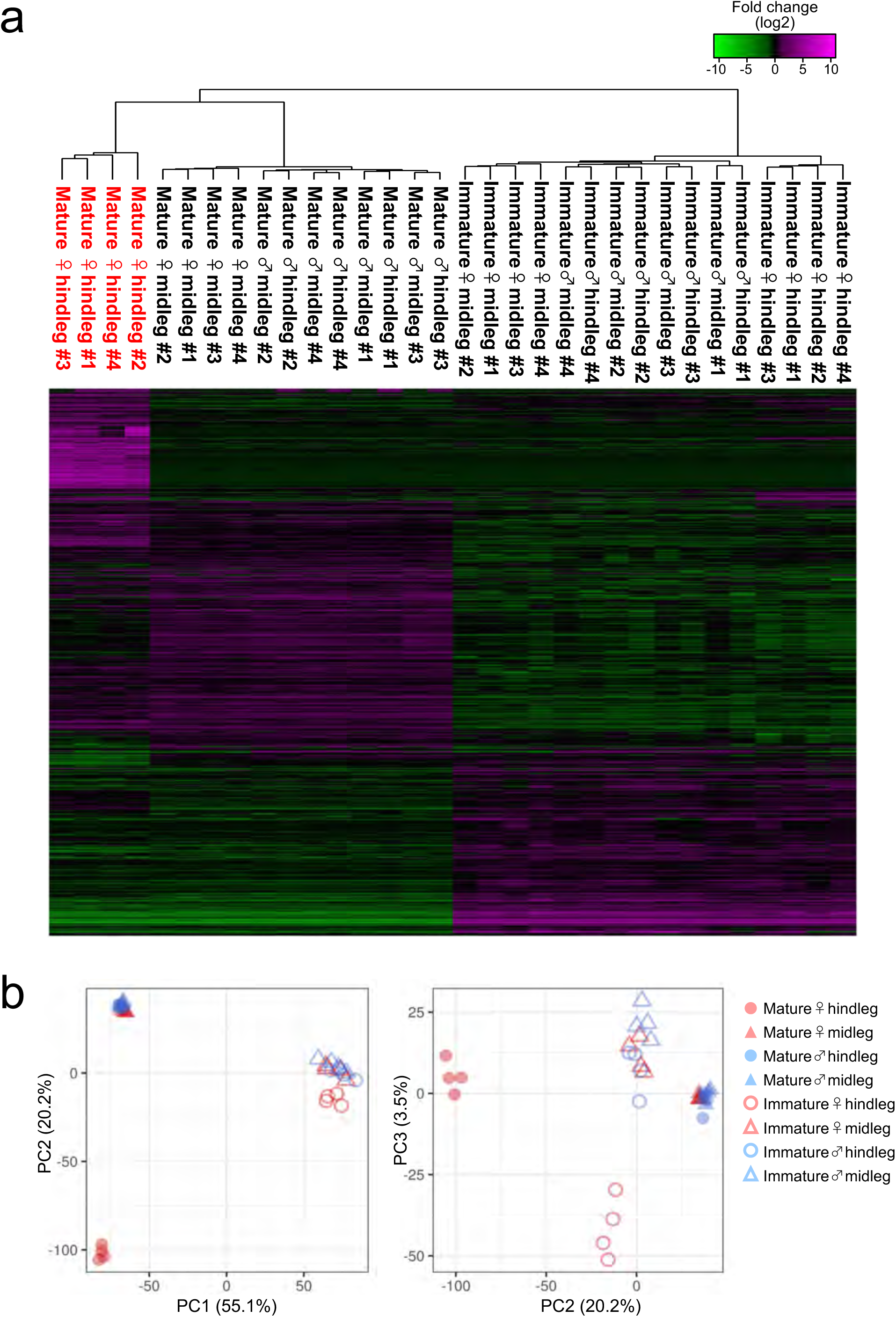
Comparative transcriptomics of hindlegs and midlegs of adult insects of *M. gracilicorne*. Using four immature males, four mature males, four immature females and four mature females, their hindlegs and midlegs were sampled and subjected to RNA sequencing analyses. (**a**) Hierarchical clustering of gene expression profiles revealed that “legs of mature adults” and “legs of immature adults” exhibit different gene expression patterns, of which “hindlegs of mature females” exhibit distinct gene expression profiles. Heatmap visualization of differential gene expression showed a set of genes are specifically up-regulated in “hindlegs of mature females”. (**b**) Principal component analysis confirmed that hindlegs of mature females exhibit distinct gene expression profiles.

**Fig. S7.**
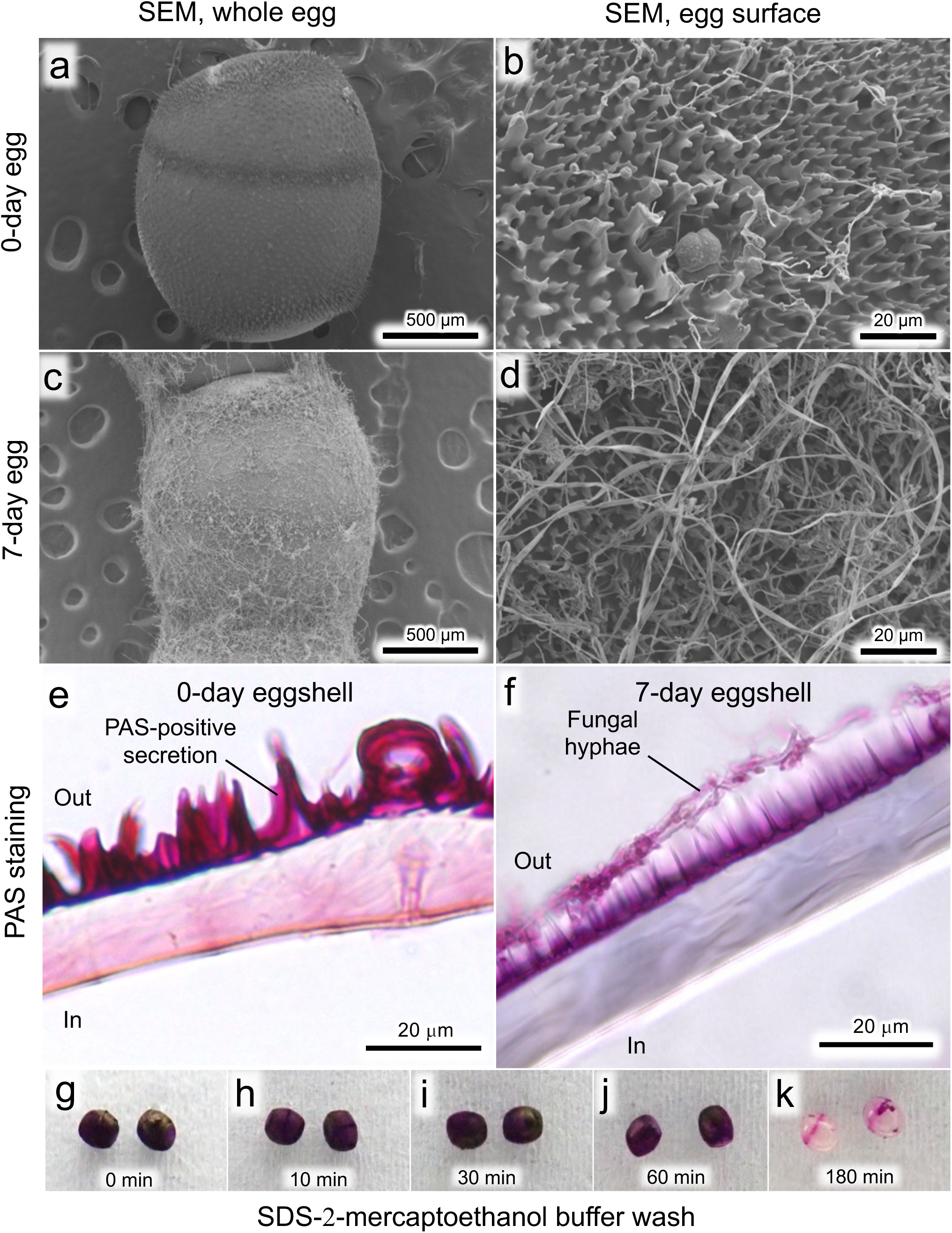
Fungal hyphae and proteinaceous secretion on egg surface of *M. gracilicorne*. (**a-d**) SEM images of egg surface. (**a, b**) 0-day egg. (**c, d**) 7-day egg. (**a, c**) Whole egg images. (**b, d**) Magnified images of egg surface. Note that the egg surface is covered with spiny secretion layer (**b**) on which fungal hyphae grow (**d**). (**e, f**) Sectioned eggshell images on which polysaccharides are visualized by PAS staining in red. (**e**) 0-day egg covered with spiny and thick PAS-positive secretion layer. (**f**) 7-day egg covered with fungal hyphae in addition to the PAS-positive secretion layer. Note that fungal hyphae never penetrate the eggshell but attach on the outer surface. (**g-k**) PAS-stained eggs after treatment with extraction buffer containing SDS and 2-mercaptoethanol for 0 min (**g**), 10 min (**h**), 30 min (**i**), 60 min (**j**) and 180 min (**k**). Note that the PAS-positive layer is extracted only in the presence of 2-mercaptoethanol and detergent (also see Fig. S8b), suggesting that the major components of the secretion layer are glycoproteins.

**Fig. S8.**
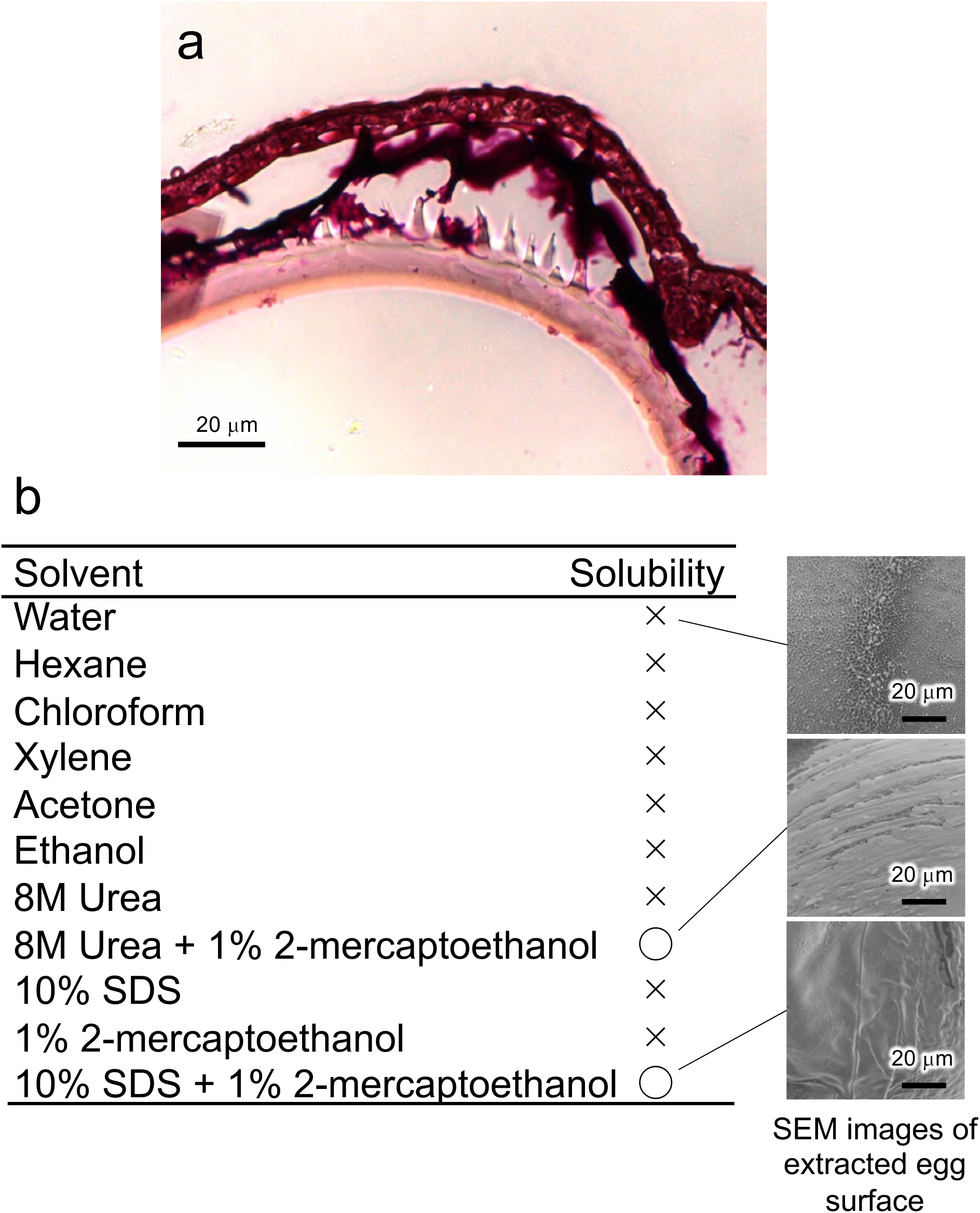
Solubility of egg surface secretion to a variety of solvents. (**a**) The spiny and thick PAS-positive secretion layer on the surface of a mature oocyte in the female’s ovary. (**b**) Solvents tested and solubility of the egg surface layer to the solvents are listed on the left side, whereas representative SEM images of the egg surface after extraction are shown on the right side. The egg surface layer is extractable when detergent and 2-mercaptoethanol coexist, suggesting that proteins crosslinked by disulfide bonds may comprise the major component of the surface layer. Considering that the surface layer is PAS-positive and thus sugar-rich, the major component of the surface layer may be glycoproteins.

**Fig. S9.**
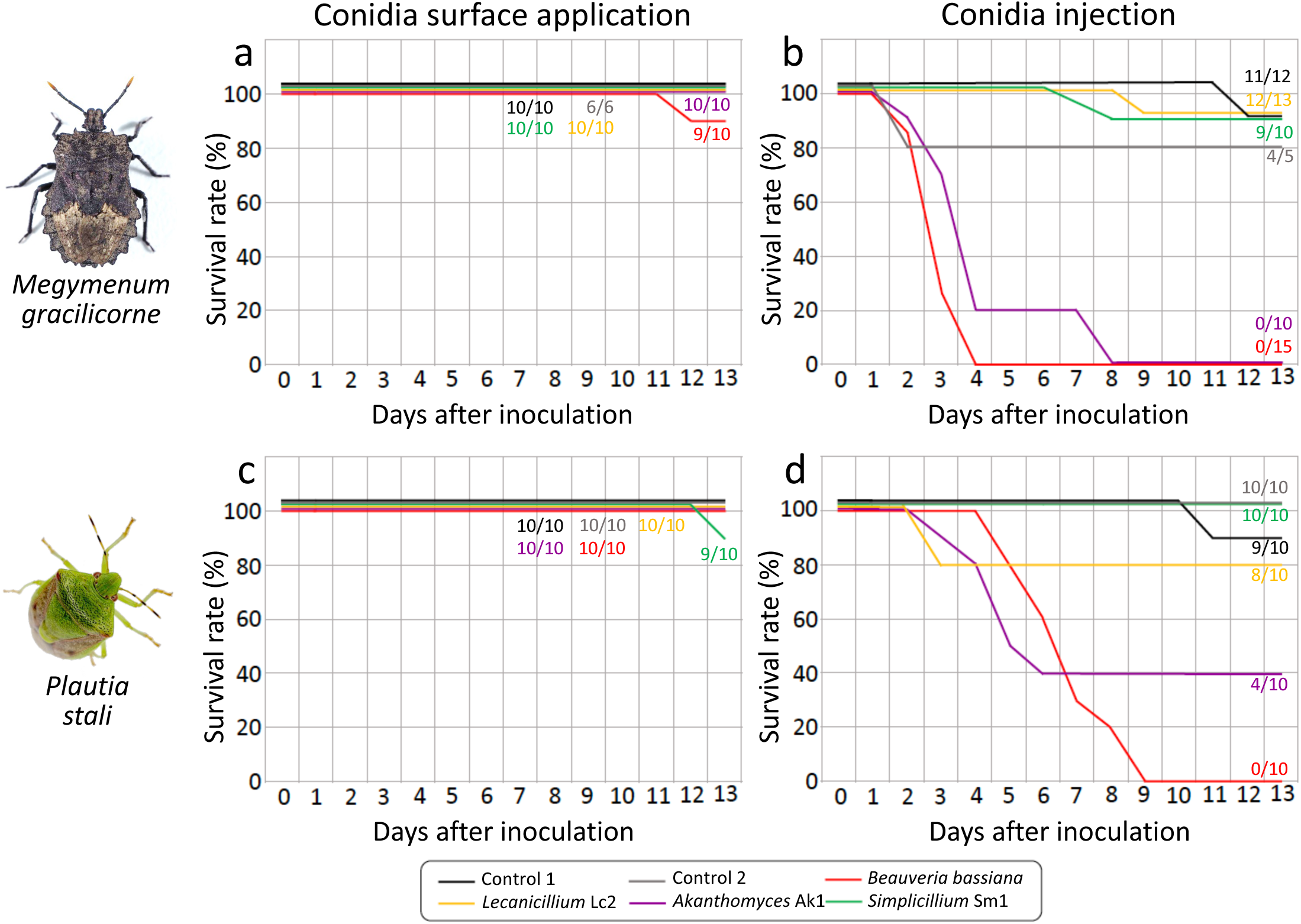
Pathogenicity of fungal strains isolated from hindleg organs of *M. gracilicorne*. (**a**) Survival curves of *M. gracilicorne* dipped in water suspending conidia of either *Simplicillium* Sm1, *Lecanicillium* Lc2, *Akanthomyces* Ak1, or *Beauveria bassiana.* For controls, the insects were dipped in water. (**b**) Survival curves of *M. gracilicorne* injected with 1.35 x 10^6^ conidia of either *Simplicillium* Sm1, *Lecanicillium* Lc2, *Akanthomyces* Ak1, or *Beauveria bassiana.* For controls, the insects were injected with water. (**c**) Survival curves of *P. stali* dipped in water suspending conidia of either *Simplicillium* Sm1, *Lecanicillium* Lc2, *Akanthomyces* Ak1, or *Beauveria bassiana.* For controls, the insects were dipped in water. (**d**) Survival curves of *P. stali* injected with 1.0 x 10^6^ conidia of either *Simplicillium* Sm1, *Lecanicillium* Lc2, *Akanthomyces* Ak1, or *Beauveria bassiana.* For controls, the insects were injected with water. Note that the conidial doses were adjusted on account of the wet body weight of the insects, 179 ± 39 mg (n = 27) for *M. gracilicorne* vs. 132 ± 21 mg (n = 37) for *P. stali*.

**Fig. S10.**
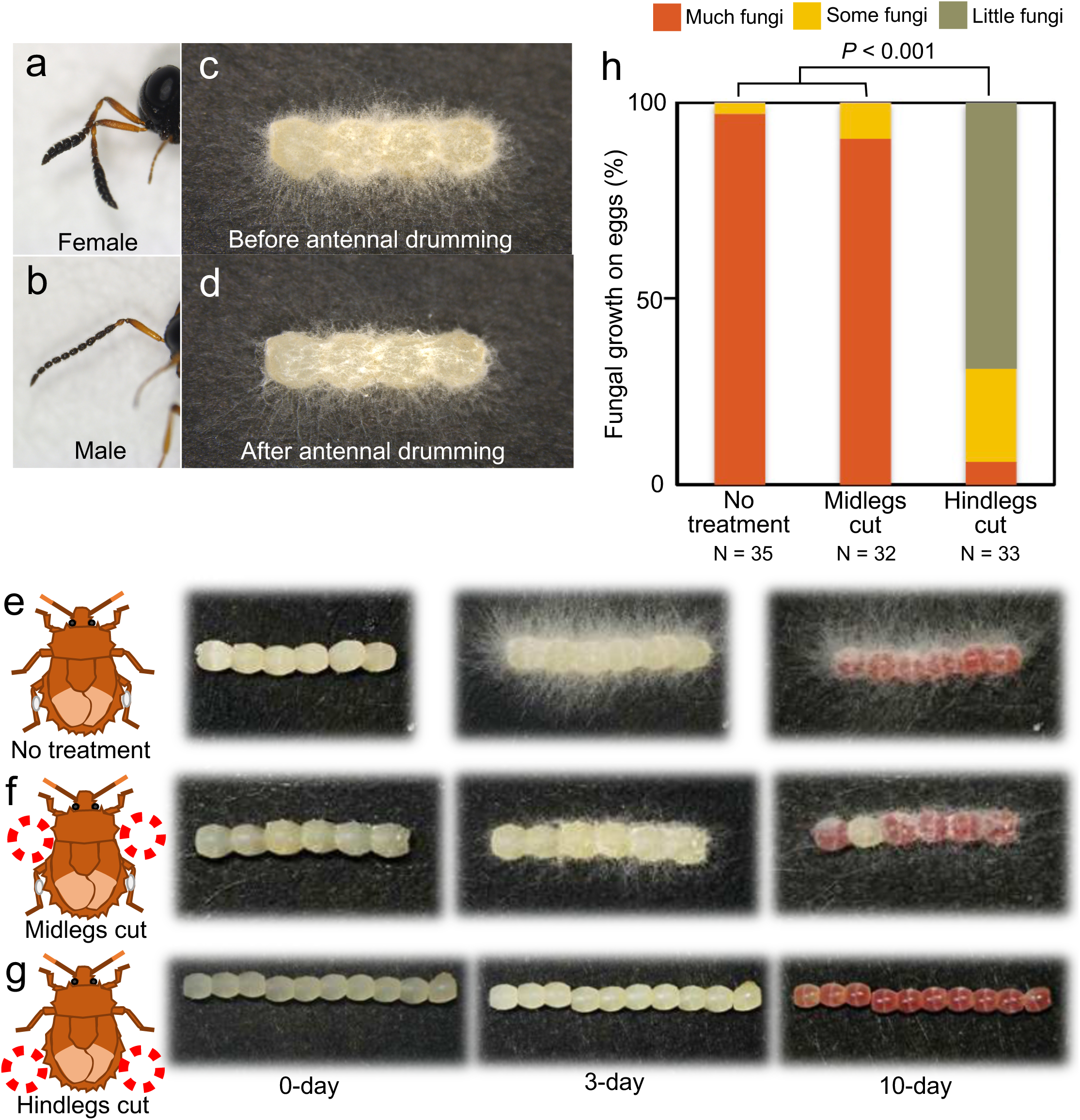
Antennae of *T. brevinotaulus*, and experimental preparation of fungus-suppressed eggs by leg amputation of reproductive adult females of *M. gracilicorne*. (**a, b**) Antennae of female (**a**) and male (**b**) of *T. brevinotaulus*. (**c, d**) A fungus-covered egg mass of *M. gracilicorne* before (**c**) and after (**d**) antennal drumming by a female of *T. brevinotaulus*. (**e-h**) Experimental preparation of fungus-suppressed eggs by leg amputation of of *M. gracilicorne*. (**e**) Eggs laid by a control adult female with no treatment. The eggs are covered with dense fungal hyphae. (**f**) Eggs laid by an adult female whose mid tibiae were experimentally amputated. With intact hindlegs, the female can transfer the fungi from the hindleg organs to the eggs upon oviposition. In this case, however, the eggs are certainly covered with fungal hyphae but not densely. (**g**) Eggs laid by an adult female whose hind tibiae were experimentally amputated. Without hindlegs, the female cannot transfer the fungi from the hindleg organs to the eggs upon oviposition. In this case, little fungal growth is seen on the eggs. (**h**) Suppressed fungal growth on eggs by amputation of hind tibiae of reproductive adult females. The levels of fungal growth on the eggs were scored as “much fungi” corresponding to the level of (**e**), “some fungi” corresponding to the level of (**f**), and “little fungi” corresponding to the level of (**g**). Statistically significant differences in the fungal growth levels were evaluated by Kruskal–Wallis test followed by Steel–Dwass test.

**Fig. S11.**
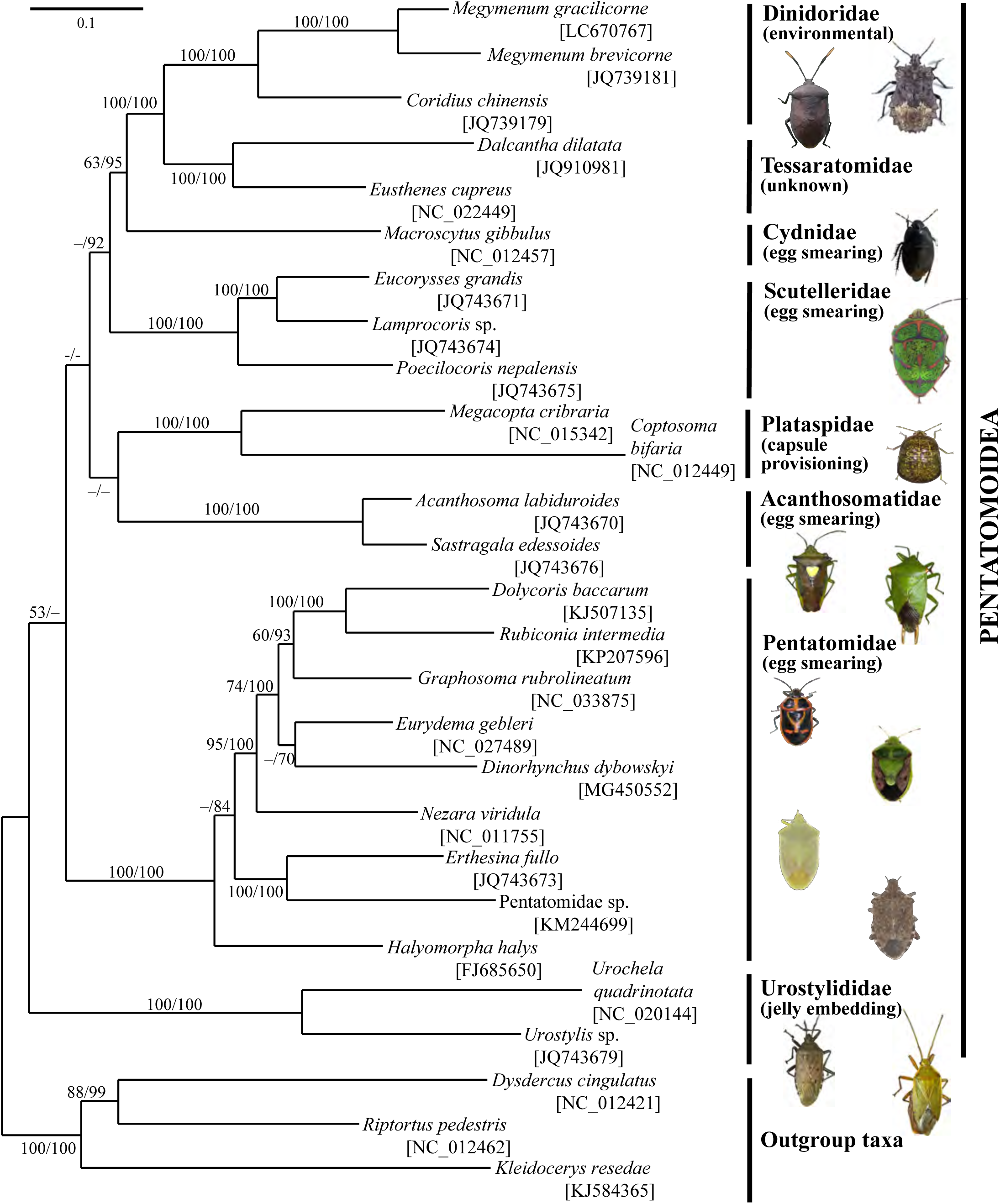
Phylogenetic placement of *M. gracilicorne* (Dinidoridae) in the superfamily Pentatomoidea. A maximum likelihood phylogeny of 24 stinkbug species representing 8 Pentatomoidea families with three outgroup taxa is inferred from amino acid sequences of 13 mitochondrial protein-coding genes consisting of 3,539 sites in total. Support values are indicated at each node in the order of bootstrap probability of maximum likelihood analysis/posterior probability of Bayesian inference. Values less than 50% are indicated as –. On the right side, Pentatomoidea family name, symbiont transmission mode in parentheses, and representative insect images are shown. The insect images are, from top to bottom, *M. gracilicorne*, *Coridius chinensis* (Dinidoriae); *Macroscytus japonensis* (Cydnidae); *Poecilocoris lewisi* (Scutelleridae); *Megacopta punctatissima* (Plataspidae); *Sastragala esakii*, *Acanthosoma labiduroides* (Acanthosomatidae); *Eurydema rugosa*, *Plautia stali*, *Nezara viridula*, *Halyomorpha halys* (Pentatomidae); *Urochela quadrinotata*, *Urostylis westwoodi* (Urostylididae). For symbiont transmission modes, see the following references: Dinidoridae (56); Scutelleridae (81); Plataspidae (82); Acanthosomatidae (83); Pentatomidae (84,85); Urostylididae (86). No information is available for Tessaratomidae. For Cydnidae, while egg smearing was reported for some species (87), other species are likely to acquire the symbiont environmentally. For Pentatomidae and Scutelleridae, both egg smearing and environmental transmission may occur (88).

#### Note S1. On fungal diversity detected from the hindleg organs and the eggs of *M. gracilicorne*

We collected reproductive adult females from different localities, each female was reared in isolation and allowed to lay two egg masses consecutively, and the hind tibiae, the first egg mass and the second egg mass were sampled and subjected to fungal cultivation to isolate up to 4 colonies per sample (Table S1; Fig. S3). In total, 631 fungal isolates were obtained and subjected to sequencing of ribosomal internal transcribed spacer (ITS) region, which yielded 572 fungal ITS sequences (with 59 unidentified isolates due to sequencing failure) (Table S1; Fig. S4). On the other hand, the samples of the hind tibiae, the first egg masses and the second egg masses were subjected to DNA extraction and ITS amplicon sequencing, which yielded 6,230,166 ITS reads. Mapping of these sequences to the fungal phylogeny revealed that (i) most sequences (92.8% of isolates; 89.9% of ITS seq reads) were placed within the class Sordariomycetes of the Ascomycota (Fig. S4), (ii) most isolates (91.6% of isolates; 83.9% of ITS seq reads) were classified to the family Cordycipitaceae mainly consisting of entomopathogenic fungi such as *Cordyceps*, *Beauveria* and others (10,11) (Fig. S5), and (iii) the majority of isolates represented several specific fungal lineages such as *Lecanicillium* (Lc1, Lc2, Lc3, etc.), *Simplicillium* (Sm1, Sm2, etc.) and *Akanthomyces* (Ak1) (12–14) (Figs. S1 and S5).

#### Note S2. On genes highly and preferentially expressed in the hindlegs of mature adult females of *M. gracilicorne*

Of 19,944 contigs classified as insect-derived genes (Table S3), we identified 550 differentially expressed genes (DEGs) that showed significantly high expression levels in the hindleg of mature females (FDR *q* < 0.001, Table S6). Functional enrichment analysis revealed an overall up-regulation of genes related to transporter activities in the hindleg of mature females (UniProt Keyword “Transport”, *P* = 2.23e^-9^). The gene category included genes for ABC transporters (25 contigs), facilitated trehalose transporter Tret1 (14 contigs), alpha-tocopherol transfer protein (3 contigs), ammonium transporter Rh type C2, and zinc transporter ZIP8. The enhanced transport activities might reflect the secretory role of the epidermal cells of the hindleg organ, where uptake and secretion of nutrients from hemolymph to mycangial pores must actively occur to foster the fungal growth.

As for individual genes, the most highly expressed DEG in the hindlegs of mature females was the gene for a Takeout family protein (TRINITY_DN2609_c0_g2_i2), and another Takeout family gene (TRINITY_DN4168_c0_g2_i1) was also highly expressed (rank 6, Table S6). Takeout family proteins are small secretion proteins that are thought to bind to and transport hydrophobic molecules (18). These proteins may represent the major carriers of hydrophobic materials secreted for the fungi growing on the hindleg organ. We also found that the hindlegs of mature females preferentially expressed UDP-glucuronosyltransferases (UGT, e.g. TRINITY_DN6402_c0_g1_i12, TRINITY_DN11711_c0_g1_i3) in addition to other glucuronate pathway genes (e.g. TRINITY_DN421_c0_g1_i3, TRINITY_DN6408_c0_g3_i2), which may be involved in enhanced mobilization of hydrophobic compounds by glucuronidation. Among the transporter genes, several genes for ABC transporters belonging to the subfamily B1, also called the multidrug resistance 1 (e.g. TRINITY_DN36_c0_g1_i19, TRINITY_DN28462_c0_g1_i1, TRINITY_DN8511_c0_g1_i2), showed remarkably specific expression in the hindleg organ (>1,000-fold increase compared to the other tissues). These transporters are known to excrete xenobiotic compounds, especially lipophilic compounds and their glucuronide conjugates (89). Therefore, in conjunction with the enhanced glucuronidation activity by UGT, these transporters may constitute a transport system that transfer some lipophilic compounds toward the cuticular surface for supporting the fungal growth.

Also, we identified a cuticular protein gene (TRINITY_DN15760_c0_g1_i1) that was dominantly expressed in the hindlegs of mature females. It may be relevant to the formation of unique cuticular structure of the hindleg organ (see Figs. 1 and 3). It should be noted that several genes constituting the Toll signaling immune pathway (90), like *spatzle* (TRINITY_DN13022_c0_g1_i2), *cactus* (TRINITY_DN229_c0_g1_i4) and *pellino* (TRINITY_DN10144_c0_g1_i4), were up-regulated in the hindlegs of mature females. The increased levels of these immune regulators are likely for defense against and/or regulation over the fungi growing on the hindleg organ.

#### Note S3. On proteins detected from the egg surface of *M. gracilicorne*

Presumably because of small quantity and simple composition of the proteins residing on the egg surface, our proteomic analysis with reference to the transcriptomic data identified only six egg surface proteins: a secretion protein with an odorant binding protein (OBP)-like motif, a β-glucuronidase-like protein, a probable antimicrobial peptide, a soma ferritin, and two uncharacterized proteins (Table S7).

The overwhelmingly abundant egg surface protein was a 16.4 kDa secretion protein with an OBP-like domain. Homology searches against the existing DNA and protein databases retrieved no closely related proteins in other organisms, except for several OBP-like proteins of other stinkbugs (91). Recently, it was reported that, in stinkbugs of the family Plataspidae, a secretion protein with an OBP-like domain, called PMDP, is mixed with symbiont cells, packaged within mother-made “symbiont capsules”, deposited with eggs upon oviposition, and essential for survival and vertical transmission of the symbiont (24). However, the egg surface protein of *M. gracilicorne* shows no phylogenetic relationship to PMDP except for the OBP-like domain. Considering the quantitative predominance, the protein might be consumed by the fungus and contribute to the hyphal growth on the egg surface, at least to some extent.

The β-glucuronidase and the probable antimicrobial peptide on the egg surface might be involved in, although speculative, defense against microbial pathogens, suppression of microbial contaminants, and/or control of the fungal proliferation on the egg surface. It may be relevant that some termites smear β-glucosidase and lysozyme on the egg surface presumably as defensive or social recognition molecules (92,93).

As for soma ferritin, its potential biological role in totally elusive. Other proteinaceous and non-proteinaceous molecules residing on the egg surface of *M. gracilicorne* should be investigated in future studies.

#### Note S4. On phylogenetic affinity of fungal strains predominantly detected from the hindleg organ of *M. gracilicorne*

Notably, previous reports generally support the low to moderate virulence of the hindleg-associated fungal strains. *L. antillanum* allied to *Lecanicillium* Lc2 and *L. aphanocladii* allied to *Lecanicillium* Lc1 (see Fig. S5) have been isolated mainly from soils, mushrooms, powdery mildews and other fungi, and also from insects. *S. subtropicum* allied to *Simplicillium* Sm1 (see Fig. S5) has been mainly isolated from soils, rust fungi, mushrooms and other fungi, and also from plants, nematodes and insects. By contrast, *A. lecanii* and *A. muscarium* (formally classified as *L. lecanii* and *L. muscarium*) allied to *Akanthomyces* A1 (see Fig. S5) have been mainly isolated from insects and fungi, and utilized as biological control agents for whiteflies, aphids, powdery mildews, etc. (12–14,25,26). It seems plausible that adult females of *M. gracilicorne* selectively establish association with the non-pathogenic fungi like *Lecanicillium* and *Simplicillium* on the hindleg organ, which entail occasional acquisitions of closely-related pathogenic fungi *Akanthomyces*, but the insects are resistant to them as long as the fungi are externally associated outside the thick cuticle.

#### Note S5. On fungal genes expressed on the hindlegs of mature adult females of *M. gracilicorne*

Of 68,154 contigs obtained from the RNA-sequencing analysis, 6,720 contigs were classified as fungus-derived genes based on BLAST sequence similarity (Table S3). Among them, 6,477 contigs (96.4%) showed predominant (>10 fold) expression in the hindlegs of mature females as compared to other tissues, and were considered as genes derived from the hindleg-associated symbiotic fungi (Table S8). Most of them (5,793 contigs, 89.4%) exhibited the best hit to protein sequences of Cordycipitaceae fungi including *Akanthomyces lecanii*, which were consistent with the results of amplicon analysis and phylogenetic analysis of the symbiotic fungi (Fig. 2; Fig S5). These contigs were clearly derived from multiple fungal species, and those with high expression levels were mainly represented by such housekeeping genes as ribosomal proteins and histone proteins. Several genes involved in hyphal growth, including actin, myosin, tubulin and SEY1 (94,95), were highly represented, which must reflect luxuriant fungal growth on the hindleg organ (see Fig. 1). We surveyed antibiotic-related genes like polyketide synthases, but no genes encoding putative polyketide synthases (type I), which consists of ketoacyl synthase, acyl transferase, and acyl carrier protein domains, etc. (96), were detected in the gene list.

**Table S1.** Samples of *M. gracilicorne* used for fungal isolation and characterization. DinidoridSym_bioRxiv_TableS1.xlsx

**Table S2.** Samples of *M. gracilicorne* used for fungal isolation and characterization. DinidoridSym_bioRxiv_TableS2.xlsx

**Table S3.** Summary of similarity-based contig assignment of RNA sequencing data of midlegs and hindlegs of immature and mature adult insects of *M. gracilicorne*. DinidoridSym_bioRxiv_TableS3.xlsx

**Table S4.** Similarity-based read assignment of RNA sequencing data of midlegs and hindlegs of immature and mature adult insects of *M. gracilicorne*. DinidoridSym_bioRxiv_TableS4.xlsx

**Table S5.** Summary of differentially expressed genes (DEGs) that were up-regulated in the hindleg organ of mature adult females of *M. gracilicorne*. DinidoridSym_bioRxiv_TableS5.xlsx

**Table S6.** List of genes preferentially expressed in the hindleg organ of mature adult females of *M. gracilicorne*. DinidoridSym_bioRxiv_TableS6.xlsx

**Table S7.** Proteins identified from egg surface of *M. gracilicorne.* DinidoridSym_bioRxiv_TableS7.xlsx

**Table S8.** List of fungal gene expression detected from the hindlegs of mature adult females of *M. gracilicorne*. DinidoridSym_bioRxiv_TableS8.xlsx

---

**Movie S1. Fungus transfer behavior from hindleg organ to egg surface by an ovipositing female of *M. gracilicorne*.** After laying each egg, the female rhythmically scratches the hyphae-covered hindleg organ with tarsal claws of the opposite hindleg, and then rubs the egg surface with the claws in a skillful manner. Also see Fig. 1m-o.

**Movie S2. Newborn nymphs hatching from fungus-covered eggs of *M. gracilicorne*.** Upon hatching, fungal hyphae attach to the body surface of the nymphs, but the fungi are subsequently lost during the nymphal development. Also see Fig. S2a.

**Movie S3. Behavior of *T. brevinotaulus* on a fungus-removed egg mass of *M. gracilicorne*.** After careful antennal drumming on the egg surface, the female wasp inserts the ovipositor into the host egg.

**Movie S4. Behavior of *T. brevinotaulus* on a densely fungus-covered egg mass of *M. gracilicorne*.** Because of the hyphal thicket, the female wasp cannot approach to the host eggs for oviposition

**Movie S5. Behavior of *T. brevinotaulus* on a sparsely fungus-covered egg mass of *M. gracilicorne*.** The female wasp presses down the hyphae by active antennal drumming, with frequently performing self-glooming behavior, and finally inserts the ovipositor into the host egg.

